# Visualizing the Epigenetic Landscape of Aging and Cellular Reprogramming: Optimized ATAC-see for Cells and Tissues

**DOI:** 10.64898/2026.07.10.737838

**Authors:** Natalie J Kirkland, Manuel A Castro, Yihao Yang, Bhargav D Sanketi, Mohammad Jaber, Veronica Lamas-Alverez, Freiya Malhotra, Juan Carlos Izpisua Belmonte, Pura Muñoz-Cánoves, Zachary A Levine

## Abstract

Spatial chromatin organization dictates cellular function and resilience, yet scalable imaging methods to quantify chromatin states in situ across aging and interventions are lacking. While ATAC-see can visualize accessible chromatin, its broader application is hindered by protocol variability, low throughput, and incompatibility with complex tissues. Here, we systematically optimize the ATAC-see workflow for robust, high-throughput quantitative imaging in fixed, adherent mammalian cells and fresh frozen tissues. We validate the platform’s sensitivity to pharmacologic remodeling and apply it to replicative, chronological, and pathological aging in primary human fibroblasts, revealing progressive age-associated chromatin opening and heterochromatin remodeling. Furthermore, we demonstrate that our optimized ATAC-see captures rapid, reversible chromatin reorganization during OSK(M)-driven partial reprogramming of aged fibroblasts. Finally, we extend a cost-effective and accessible protocol to murine tissue sections, quantifying *in situ* age-dependent remodeling. This standardized framework establishes chromatin accessibility as a highly scalable, sequencing-compatible imaging biomarker for evaluating aging and rejuvenation.

**Summary Statement:** ATAC-see was optimized for scalable, quantitative imaging of chromatin remodeling during aging and cellular reprogramming, and extended to characterize age-associated epigenetic changes across organs.

## Introduction

Aging is accompanied by pervasive changes in chromatin organization that affect transcriptional programs, genome stability, and cellular identity (López-Otín et al., 2013; Sen et al., 2016). These alterations include global shifts in chromatin state via redistribution or loss of facultative and constitutive heterochromatin, and remodeling of histone modification landscapes (Benayoun et al., 2019; Cecco et al., 2013; Shah et al., 2013; Zhang et al., 2015). The resulting changes in chromatin accessibility, in particular, have emerged as a robust functional readout of regulatory potential across cell types and age (Buenrostro et al., 2013; Cusanovich et al., 2018; Lu et al., 2026; Ucar et al., 2017).

Genome-wide assays such as DNase-seq and ATAC-seq have revolutionized the mapping of accessible chromatin, enabling locus-resolved measurements of regulatory potential in development, homeostasis, and aging (Benayoun et al., 2019; Buenrostro et al., 2013; Lu et al., 2026; Ucar et al., 2017). These sequencing-based studies have shown that aging is accompanied by extensive, region-specific changes in accessibility in multiple tissues, often with net increases at regulatory and repeat-rich regions and focal losses at lineage-defining and stress-response loci (Benayoun et al., 2019; Lu et al., 2026; Ucar et al., 2017). Parallel work has demonstrated progressive loss of heterochromatin, lamina detachment, and aberrant opening of normally repressed regions in aging, including in models of accelerated aging such as Hutchinson Gilford progeria syndrome (HGPS) and Werner syndrome (Cecco et al., 2013; Liu et al., 2011; Shah et al., 2013; Shumaker et al., 2006; Yi and Kim, 2020; Zhang et al., 2015). However, sequencing-based accessibility assays do not preserve spatial context and have relatively low throughput in terms of the number of conditions, time points, or donors that can be affordably surveyed in a single experiment. They also generally require fresh or nuclei-compatible material and are not readily applicable to archived fixed samples or cryosections, which are common in aging cohorts and biobanks.

Microscopy-based approaches that directly visualize accessible chromatin in intact cells and tissues would therefore be highly valuable, as they would enable quantitative interrogation of chromatin state in relation to nuclear architecture, cell morphology, and tissue anatomy and are amenable to high-throughput imaging and quantitative analysis. Among these, ATAC-see was developed as a microscopy adaptation of ATAC-seq that uses Tn5 transposase loaded with fluorescently-labelled oligonucleotides to label open chromatin in cells (Chen et al., 2016; Xie et al., 2020). Early implementations demonstrated that ATAC-see can reveal nuclear regions enriched for open chromatin and can be combined with immunostaining for histone modifications or DNA dyes, suggesting potential for multiplexed epigenetic imaging (Chen et al., 2016; Xie et al., 2020).

Nevertheless, several technical limitations have restricted ATAC-see’s broader use. First, the precise fixation, permeabilization, and tagmentation parameters have not yet been systematically optimized to achieve the signal-to-noise ratios necessary for large, image-based screens. For instance, a recent genome-wide application of ATAC-see revealed that even low levels of fixation distorted the expected intensities of euchromatin relative to heterochromatin when compared to DAPI staining (Ishii et al., 2024). However, this study used only a 10-min tagmentation step, which is likely insufficient for the large Tn5 transposase complex to fully diffuse through a crosslinked nucleus, highlighting the critical need for rigorous protocol calibration. Second, the original ATAC-see and 3D ATAC-PALM protocols were designed primarily for cultured cells; and there are still no robust protocols for complex tissue architecture, particularly fresh frozen sections. Third, the technique has suffered from inconsistent performance in different laboratories, often characterized by low tagmentation efficiency, high background signal, and unexpected findings in disease models such as HGPS (Köhler et al., 2020).

Beyond resolving these technical barriers, a key motivation for optimizing ATAC-see is to enable the cost-effective assessment of chromatin states across large arrays of conditions, compounds, and interventions without immediate recourse to sequencing. ATAC-seq and related genomic assays are expensive, labor-intensive, and difficult to scale for early-stage screening. In contrast, ATAC-see yields a global, nucleus-wide readout of chromatin accessibility at single-cell resolution and can be implemented on standard microscopy platforms. This enables rapid prioritization of conditions that elicit robust chromatin responses prior to in-depth genomic profiling. Importantly, ATAC-see reports on total accessible chromatin, integrating signals across many regulatory elements and nuclear compartments. This provides a holistic view of nuclear architecture that is distinct from immunofluorescence staining of specific histone marks (e.g., H3K9me3, H3K27ac), which detects individual chromatin features but does not directly measure overall physical accessibility. Furthermore, an optimized ATAC-see protocol can be performed in under three hours, less than half the time required for standard immunofluorescence. An efficient and scalable assay is particularly desirable in the aging field, where the search for interventions that reverse epigenetic decline is rapidly expanding. For example, transient expression of the Yamanaka factors (Oct4, Sox2, Klf4, c-Myc: herein, OSKM) can rejuvenate multiple epigenetic features without full loss of cellular identity (Ocampo et al., 2016; Olova et al., 2019; Sarkar et al., 2020).

Here, we comprehensively optimize and extend the ATAC-see protocol for fixed cells and tissues in the context of aging. We first perform a systematic parameter exploration of fixation and tagmentation conditions to maximize the ATAC-see signal-to-noise ratio while preserving nuclear morphology and compatibility with multiplexed histone immunofluorescence. We then demonstrate that this imaging framework successfully generates high-quality ATAC-seq profiles downstream, linking spatial accessibility with genomic analysis. We demonstrate that optimized ATAC-see sensitively detects trichostatin A (TSA)-induced chromatin opening in primary human dermal fibroblasts and yields coherent changes in accessible-chromatin foci and histone modification profiles. We then apply this framework to replicative aging and to primary dermal fibroblasts from donors spanning adult and pathological aging, including HGPS, and observe progressive, age-associated increases in chromatin accessibility and heterochromatin remodeling. We subsequently use ATAC-see to visualize dynamic chromatin remodeling during OSK(M)-based continuous and partial reprogramming of aged fibroblasts, revealing multiphase epigenetic rewiring of OSKM reprogramming and recovery of chromatin landscapes in transient OSKM expression. Finally, we develop and validate a protocol that extends ATAC-see to fresh frozen murine tissue sections, enabling *in situ* quantification of age-dependent chromatin remodeling in skin, liver, and heart.

Together, our work establishes a unified, standardized ATAC-see platform spanning fixed cells, reprogramming paradigms, primary human fibroblasts, and murine tissues. By demonstrating sensitivity to pharmacologic and genetic perturbations, as well as to chronological and pathological aging, we propose ATAC-see derived chromatin accessibility metrics as candidate imaging biomarkers for aging and rejuvenation that are well suited for high-throughput, cost-efficient experimental and translational applications.

## Results

### Standardization of ATACsee for fixed cells enables high-throughput imaging and quantification

To develop a robust and scalable ATAC-see workflow, we recognized that short-term storage of large sample cohorts, together with strict preservation of cellular and nuclear morphology for accurate spatial imaging, are required. However, previous literature indicates that introducing even low levels of fixation severely diminishes the ATAC-see intensity and distorts the expected chromatin readouts (Ishii et al., 2024; Köhler et al., 2020). We reasoned that this fixation-induced signal loss may occur because the crosslinked nuclear matrix acts as a physical diffusion barrier to the bulky Tn5 transposase complex during standard, short tagmentation windows. To overcome the inconsistent tagmentation efficiency and high background signal that have limited ATAC-see, we systematically examined the impact of varying fixation and permeabilization conditions paired with extended tagmentation durations in primary human dermal fibroblasts (HDFs).

To perform the assay, cells cultured in 96-well plates were treated with in-house expressed and purified Tn5 transposase, which was pre-loaded with Cy5-labeled Mosaic End (ME) DNA oligonucleotide adapters as previously described (Chen et al., 2016). Following treatment, the cells were imaged using a Nikon Ti2 inverted widefield microscope. To create a standardized analysis workflow (see Materials and Methods), images were processed using Nikon Elements, CellPose, and Arivis software. We extracted nuclear morphology and ATAC-see signal intensity, and measured background Tn5 binding (using EDTA-inhibited controls), to identify an optimal parameter window across a multidimensional matrix.

Using 1% PFA fixation for 10 min, we observed minimal variation in nuclear area and circularity following ATAC-see (Fig. 1A, B; Fig. S1B-D). Importantly, tagmentation times of 30 min or less yielded little nuclear intensity above the EDTA-treated background; in fact, 10-min tagmentation resulted in signal below background in samples fixed with >0.5% PFA (Fig. 1C, E; Fig. S1A, D). Conversely, extended tagmentation (120 or 240 min) generated increasing signal even at higher fixation, suggesting non-biological, over-tagementation of cleaved DNA fragments entrapped within the dense crosslinked matrix (Fig. 1C, E; Fig. S1A, D).

**Figure 1.**
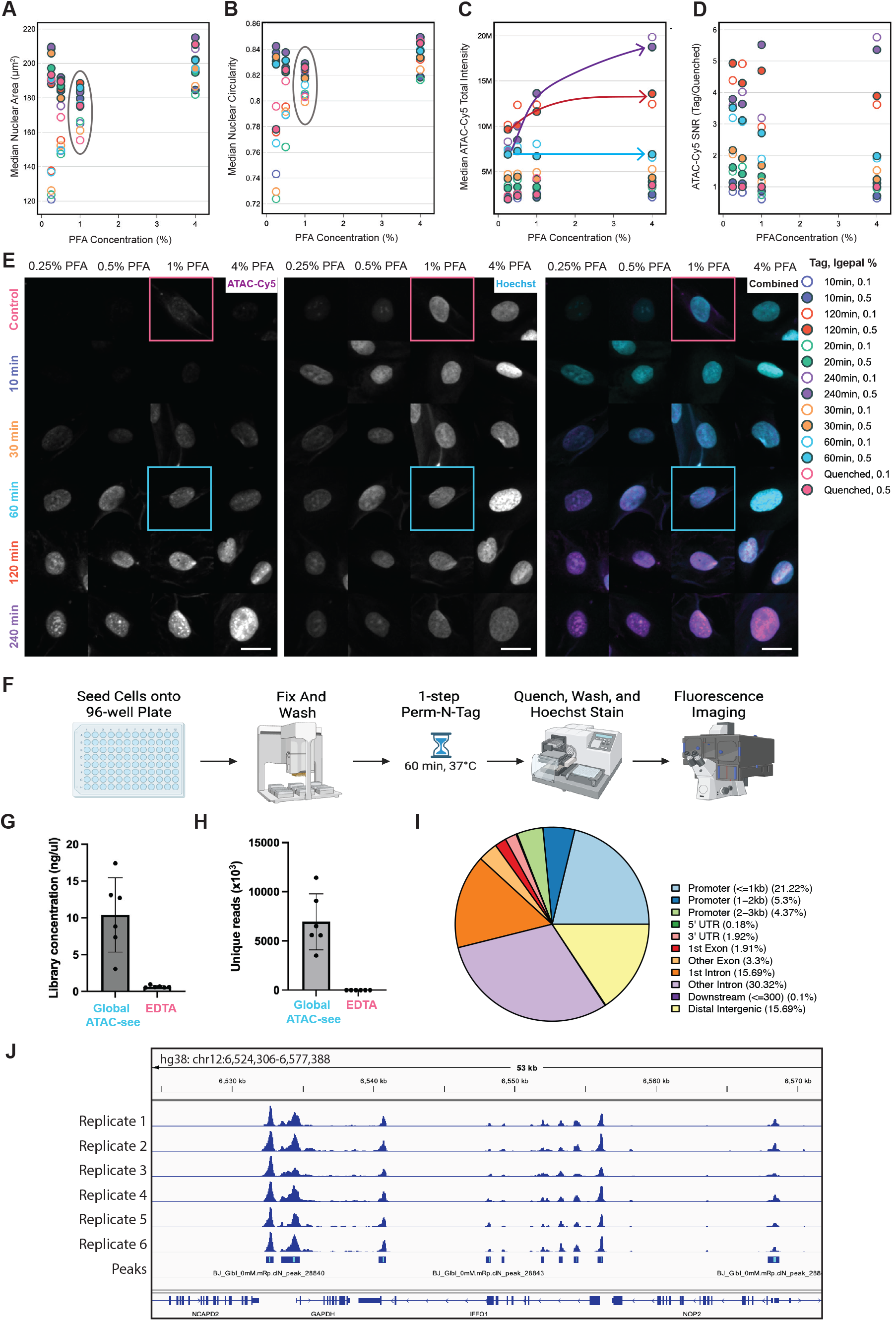
Optimization of *in situ* ATAC-see fixation and tagmentation parameters in primary fibroblasts. (A–B) Impact of formaldehyde (PFA) fixation concentration on nuclear morphology, quantified by median nuclear area (A) and median nuclear circularity (B). Data points represent the median values of distinct experimental wells, color-coded by tagmentation time and permeabilization condition (Igepal concentration). (C–D) Quantitative assessment of ATAC-see assay performance across varying PFA concentrations. Median ATAC-Cy5 total intensity (C) and the signal-to-noise ratio (SNR; calculated as the ratio of active tagmentation signal to EDTA-quenched background) (D) are shown. Arrows in (C) highlight the distinct trajectories of total intensity at varying tagmentation times. (A-D) Data shown are representative from a single 96-well plate experiment, which was performed independently twice (*N* = 2 biological replicates). A minimum of *n* > 1,000 single nuclei were analyzed per condition. (E) Representative high-throughput widefield images illustrating the influence of fixation stringency and tagmentation duration. ATAC-Cy5 signal (magenta) tracks chromatin accessibility, while Hoechst (cyan) defines total nuclear DNA. Scale bars equal 20µm. Boxes refer to the conditions sequenced in (G-J). (F) Schematic of rapid ATAC-see high-throughput imaging protocol. (G-J) Global ATAC-see yields high-quality, genome-wide chromatin accessibility profiles in BJ fibroblasts (G) Sequencing library concentration (ng/µl) recovered from Global ATAC-see versus the EDTA-treated control. Bars represent the mean and s.d. (n = 6 independent replicates). (H) Number of unique (non-duplicate) aligned reads (×10³) per condition, plotted as in (G). (I) Genomic distribution of accessible chromatin regions, annotated relative to gene features using ChIPseeker. Annotation was performed on the pooled peak set across all replicates. Peaks are consistent with the expected genome-wide accessibility landscape. (J) Representative genome-browser view (IGV) of Global ATAC-see signal at the *GAPDH* locus and surrounding genes (hg38, chr12:6,524,306–6,577,388; 53 kb window). Tracks show read coverage for the six BJ neonatal dermal fibroblasts with the pooled peak set shown below ("Peaks"). Accessibility peaks are highly reproducible across all replicates and localize to promoters and gene bodies.

Ultimately, 1% PFA fixation combined with a 60-min tagmentation provided a robust signal-to-noise ratio, preserving the structural integrity necessary for sample storage without introducing artificial signals. This window also maintained compatibility with follow-up multiplexed immunofluorescence, which often requires increased detergent concentrations for permeabilization (Fig. 1A, B, D, E; Fig. S1B-D). Furthermore, ATAC-see signal intensity positively correlated with DNA content, providing a normalization baseline to account for cell cycle differences in downstream studies. (Fig. S1E). The optimized tagmentation conditions also proved robust to increased fixation time (18 min) and lower cell density (3,000 cells/cm² vs. 15,000 cells/cm²) (Fig. S1F). By integrating this workflow with automated plate washers and liquid handlers, and high-throughput analysis of thousands of nuclei to derive our nuclear metrics, we established a standardized, quantitative, and highly scalable ATAC-see pipeline for fixed cells (Fig. 1F).

To verify that the fixation and extended tagmentation steps remained compatible with downstream sequencing, cells cultured and imaged in glass-bottom 12-well plates were subsequently reverse-crosslinked. The resulting library concentrations were significantly greater than those of EDTA-treated negative controls, yielding an average of 7 million unique reads across six replicates (Fig. 1G, H). The near-absence of recoverable libraries and unique reads in the EDTA controls confirmed that the sequencing signal is strictly tagmentation-dependent. Analysis of the genomic distribution of accessible chromatin regions, annotated relative to gene features using ChIPseeker, revealed an enrichment of peaks at promoter-proximal regions (≤1 kb, 21.22%) (Fig. 1I). Peaks were also broadly distributed across intronic (first intron, 15.69%; other introns, 30.32%) and distal intergenic (15.69%) regions, consistent with expected genome-wide accessibility landscapes (Yan et al., 2020). Furthermore, accessibility peaks were highly reproducible across all replicates and correctly localized to promoters and gene bodies, as demonstrated by visualization of the *GAPDH* locus (Fig. 1J). Collectively, these data establish that quantitative imaging can be successfully coupled with sequencing, thereby broadening the utility of the optimized ATAC-see pipeline.

### ATAC-see detects TSA-induced chromatin opening

To validate the sensitivity of our optimized ATAC-see platform for detecting dynamic chromatin remodeling, we treated HDFs with increasing concentrations of the histone deacetylase (HDAC) inhibitor TSA (62.5 nM–1 µM) for 4 h (Fig. 2C). While the raw ATAC-Cy5 (ATAC-see) sum intensity increased upon TSA treatment (Fig. S2A), we normalized these values to DNA content to account for TSA-induced nuclear expansion and cell cycle variations. This normalized metric confirmed a significant, dose-dependent increase in global chromatin accessibility (Fig. 2A, B; Fig. S2B-D).

**Figure 2.**
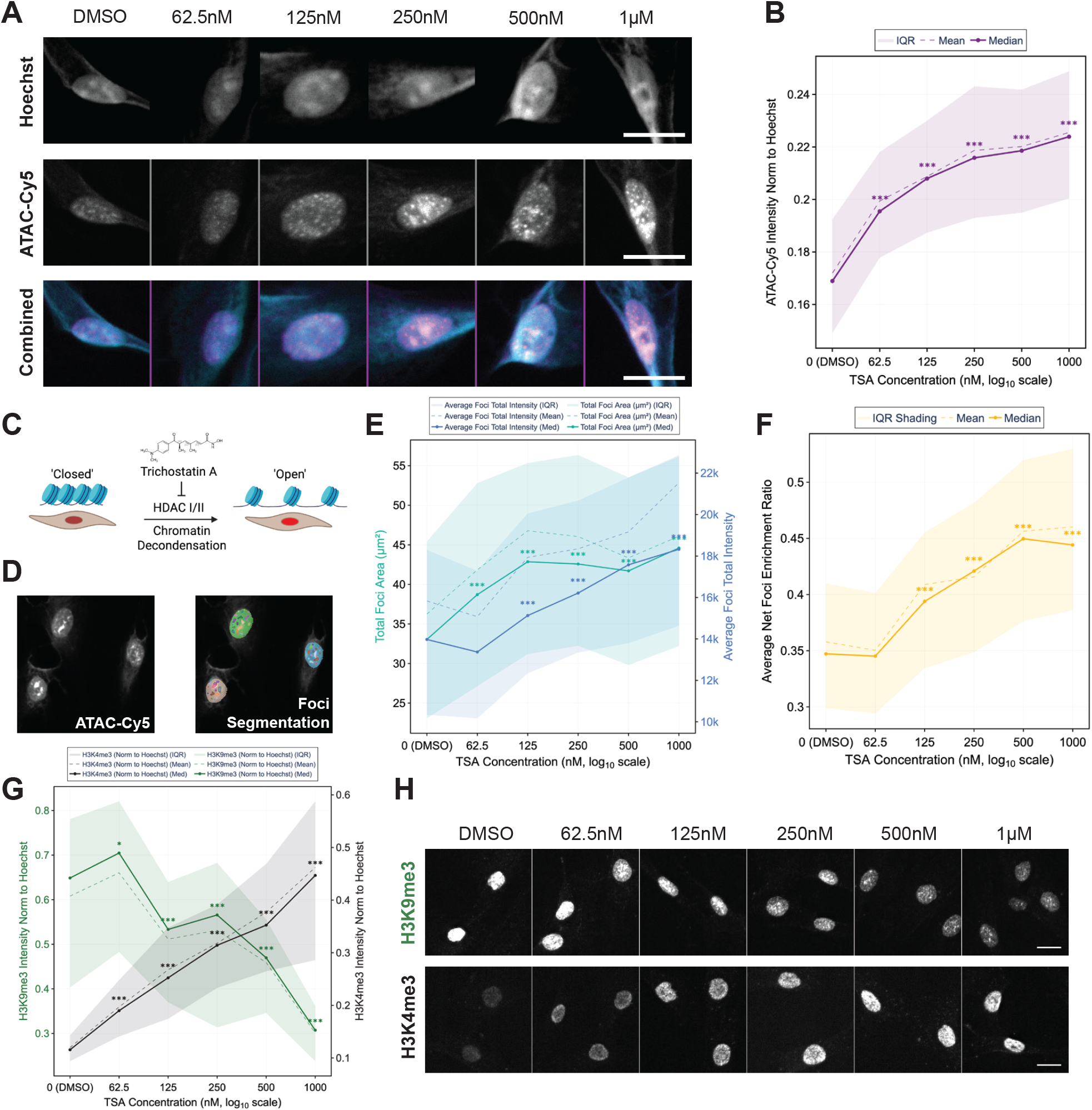
*In situ* ATAC-see accurately quantifies TSA-induced global chromatin decondensation and spatial reorganization. (A) Representative widefield images of 75-year-old primary human dermal fibroblasts (GM08401) treated with a dose-response gradient of TSA (DMSO, 62.5 nM, 125 nM, 250 nM, 500 nM, 1 µM). Top to bottom: Hoechst, ATAC-Cy5, and composite. Scale bars, 20 µm. (B) Line graph quantifying global chromatin accessibility, represented by ATAC-Cy5 intensity normalized to total nuclear Hoechst signal across the TSA dose-response. (C) Schematic overview of trichostatin A (TSA) treatment, an HDAC I/II inhibitor that drives structural transition from a compact ("closed") to a highly accessible ("open") chromatin state. (D) Representative image demonstrating the automated spatial foci segmentation mask applied to the ATAC-Cy5 channel. (E, F) Quantification of localized spatial remodeling upon TSA-induced decondensation with line graphs displaying a dual-axis overlay of total foci area per nucleus (µm²) (turquoise) and average foci intensity (blue) (E) and average net foci enrichment (F). (A-F) Data shown are representative from a single 96-well plate experiment, which was performed independently more than 3 times (N ≥ 3 biological replicates). A minimum of n ≥ 272 single nuclei were analyzed per condition across 6 well replicates. (G) Parallel immunofluorescence quantification presented as dual-axis line graph for the constitutive heterochromatin mark H3K9me3 (green) and the active euchromatin mark H3K4me3 (black), normalized to Hoechst intensity, validating the biochemical transition accompanying physical decondensation. For all line graphs, solid and dashed lines represent the median and mean values, respectively, while the shaded regions represent the interquartile ranges. Data are representative of a single 96-well plate experiment, which was performed independently more than 3 times (N ≥ 3 biological replicates). A minimum of n ≥ 207 single nuclei were analyzed per condition. (H) Representative single-channel images for H3K9me3 and H3K4me3. Statistical significance across treatment groups: *** P < 0.001, ** P < 0.01; two-sided Mann-Whitney U tests with Bonferroni correction.

As a primary advantage of ATAC-see over sequencing-based assays is the preservation of spatial context, we next investigated the sub-nuclear distribution of newly accessible chromatin using automated foci analysis. We observed that individual and total foci areas, and foci intensity exhibited striking dose-dependent increases following TSA treatment (Fig. 2E; Fig. S2E-I). Furthermore, analysis of foci intensity relative to total nuclear intensity (the foci net enrichment ratio) revealed that localized accessibility likely drives the overall nuclear signal increase (Fig. 2F).

To orthogonally confirm that these physical shifts in accessibility corresponded to expected biochemical epigenetic remodeling, we evaluated canonical histone marks in parallel. Consistent with the ATAC-see readouts, immunofluorescence showed a concurrent loss of the heterochromatin mark H3K9me3 and a gain of the active euchromatin mark H3K4me3 following TSA treatment (Fig. 2G, H; Fig. S2J-L). Together, these results demonstrate that our optimized ATAC-see pipeline is highly sensitive to TSA- induced chromatin opening, providing a robust spatial framework for profiling epigenetic drug responses and complex chromatin state transitions.

### Replicative and chronological aging are associated with increased chromatin accessibility

We next investigated whether ATAC-see can detect age-related chromatin remodeling in HDFs, beginning with a model of replicative aging across low, middle, and high passages. Using our optimized protocol, we found that DNA-normalized ATAC-Cy5 intensity increased progressively with passaging (Fig. 3A, B). This revealed a global gain in chromatin accessibility in late-passage cells that corresponded with the expected morphological expansion of nuclear area (Fig. S3A-D) (Cecco et al., 2013; Shah et al., 2013). Importantly, this stepwise increase in accessibility and nuclear expansion was preserved across alternative fixation methods, confirming the robustness of the readout (Fig. S3E, F).

**Figure 3.**
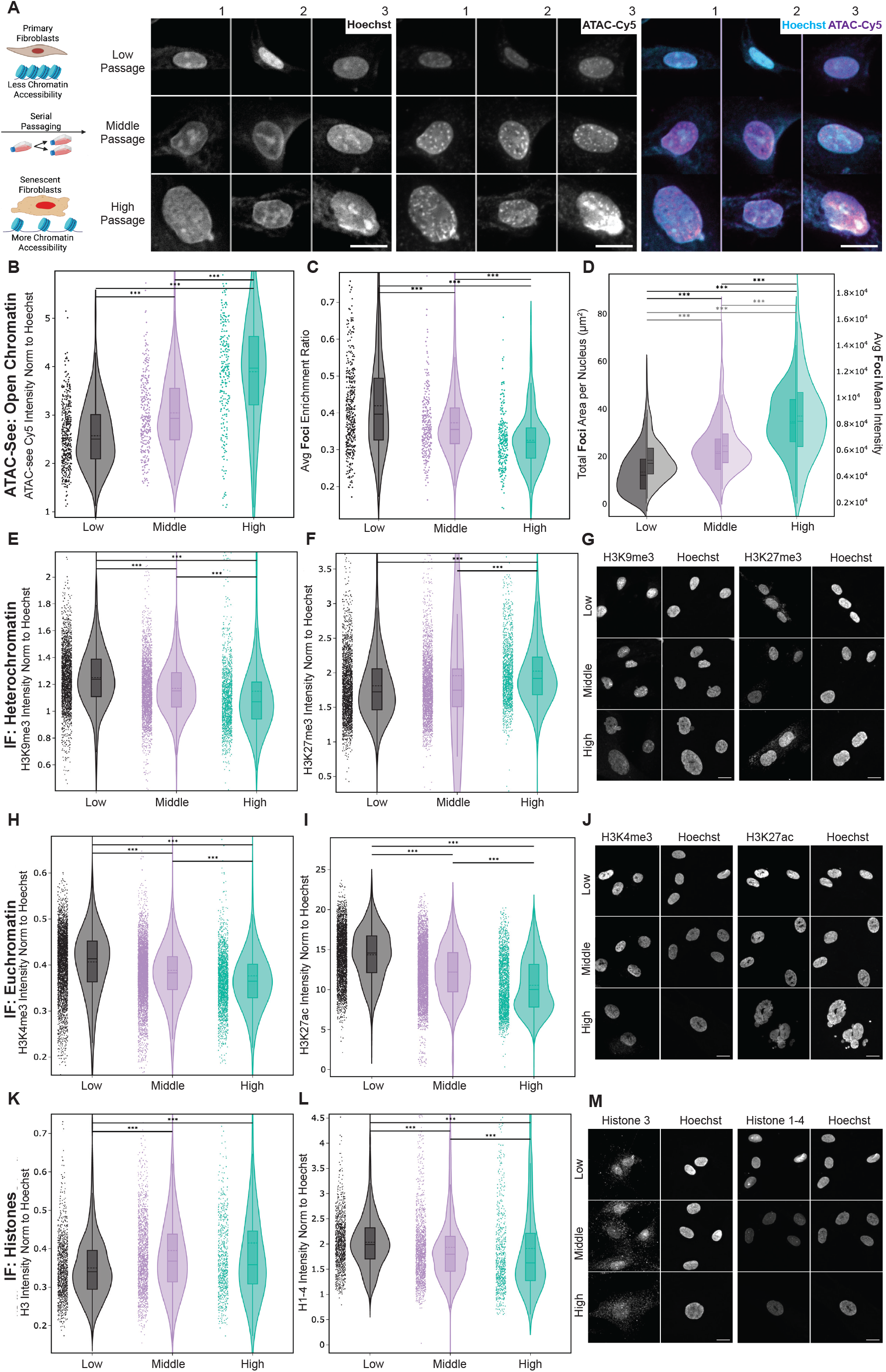
Continuous *in vitro* aging drives global chromatin decondensation, loss of heterochromatin, and structural remodeling in primary human fibroblasts. (A) Schematic overview of the serial passaging model, highlighting the transition from a compact, low-passage state to an open, high-passage state. Three representative nuclei with Hoechst (cyan) and ATAC-Cy5 (magenta) across low, middle, and high passage populations. Scale bars, 20µm. (B–D) Quantitative single-nuclei ATAC-see metrics tracking chromatin accessibility and spatial reorganization during passaging, including ATAC-see Cy5 Intensity normalized to Hoechst (B), Average Foci Net Enrichment Ratio (C), and Total Foci Area per Nucleus (left-axis) with Average Foci Mean Intensity (right-axis) (D). Data are representative of a single 96-well plate experiment, which was performed independently more than 3 times (N ≥ 3 biological replicates). n = 376; 280; 243 single nuclei for low, medium and high passage respectively across 3 well replicates. (E–G) Immunofluorescence quantification of constitutive and facultative heterochromatin. Violin plots display H3K9me3 (E) and H3K27me3 (F) intensities normalized to Hoechst, with corresponding representative widefield images (G). (H–J) Immunofluorescence of active euchromatin marks, detailing H3K4me3 (H) and H3K27ac (I) normalized intensities, accompanied by representative images (J). (K–M) Immunofluorescence of core Histone H3 (K) and pan-Histone marker, H1-4 (L), alongside representative images (M). Data are representative of a single 96-well plate experiment, n ≥ 1015; 1471; 639 single nuclei for low, medium and high passage respectively across 3 well replicates. Statistical significance across passage groups Statistical significance across treatment groups: *** P < 0.001, ** P < 0.01; * P < 0.05; two-sided Mann-Whitney U tests with Bonferroni correction.

Spatial analysis further revealed that passaging increased the count, area, and absolute mean intensity of individual accessible foci but decreased the overall foci net enrichment ratio (Fig. 3C, D and Fig. S3G-J). Therefore, this suggests that the widespread increase in chromatin accessibility dominates the increase in total nuclear fluorescence, resulting in reduced spatial contrast. This suggests that increased global chromatin accessibility during replicative aging is driven by canonical heterochromatin loss, rather than a few discrete hubs (Cecco et al., 2013; Tsurumi and Li, 2012; Villeponteau, 1997; Zhang et al., 2015). Concordantly, parallel immunofluorescence profiling of the constitutive heterochromatin mark H3K9me3 exhibited a passage-dependent decline that inversely correlated with global ATAC-Cy5 intensity (Fig. 3E, G and Fig. S3K, L). In contrast, the facultative heterochromatin mark H3K27me3 exhibited inconsistent dynamics, with a trend towards increased abundance with passage (Fig. 3F, G and Fig. S3M). Despite the global increase in chromatin accessibility, levels of the active euchromatin marks H3K4me3 and H3K27ac progressively declined across passages (Fig. 3H-J and Fig. S3N, O). This decay of canonical active marks was accompanied by a general depletion of core and linker histones H1–H4, even though total histone H3 moderately increased, reflecting large-scale nucleosomal remodeling (Fig. 3K-M and Fig. S3P, Q). The simultaneous erosion of both repressive and activating epigenetic signatures indicates a flattening of the epigenetic landscape, which is reflected in the ATAC-see measurements (Cecco et al., 2013; O’Sullivan et al., 2010).

To extend these findings to chronological and pathological aging, we profiled primary dermal fibroblasts from human donors aged 22, 75, and 96 years, alongside fibroblasts from a 14-year-old with HGPS (Fig. 4A). Although cell lines were analyzed at the lowest, most closely matched passages possible, exact passage matching across diverse donor lines remains inherently challenging. However, consistent with our replicative aging model, DNA-normalized ATAC-Cy5 intensity progressively increased with donor age and was significantly elevated in the HGPS fibroblasts, indicating a severe, disease-associated expansion of chromatin (Fig. 4A, B; Fig. S4A-C).

**Figure 4.**
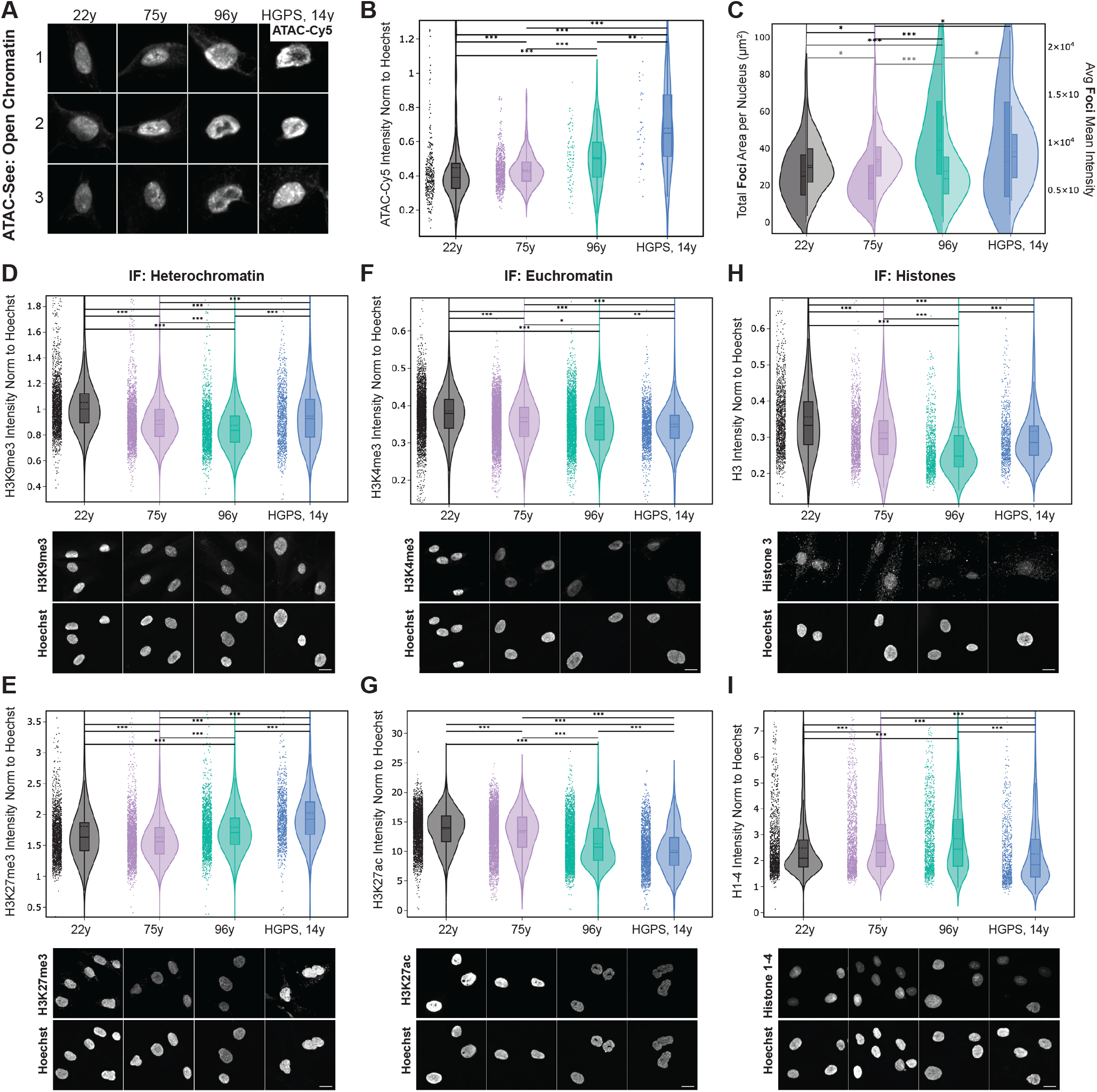
Physiological aging and Hutchinson-Gilford Progeria Syndrome drive global chromatin decondensation and epigenetic erosion in primary human fibroblasts. (A) Three representative nuclei stained with ATAC-Cy5, illustrating the progressive loss of spatial chromatin organization across a chronological aging cohort, comprising donors aged 22y, 75y, and 96y, compared to a 14-year-old patient with Hutchinson-Gilford Progeria Syndrome. Scale bars, 20µm. (B–C) Quantitative single-nuclei ATAC-see metrics assessing global chromatin accessibility and localized structural remodeling, including ATAC-Cy5 Intensity normalized to Hoechst (B) and Total Foci Area per Nucleus and Average Foci Mean Intensity (C). Data are representative of a single 96-well plate experiment, which was performed independently more than 3 times (N ≥ 3 biological replicates). n ≥ 305; 704; 75; 36 single nuclei for 22y, 75y, 96y and HGPS cell lines respectively across 2 well replicates. (D–E) Parallel immunofluorescence quantification of constitutive and facultative heterochromatin, displaying H3K9me3 (D) and H3K27me3 (E) intensities normalized to Hoechst, with corresponding representative images. (F–G) Parallel quantification of active euchromatin marks, detailing normalized intensities for H3K4me3 (F) and H3K27ac (G), accompanied by representative images. (H–I) Assessment of structural nucleosome components during aging, showing normalized intensities for core Histone H3 (H) and pan-histones marker, H1-4 (I). Data are representative of a single 96-well plate experiment, n ≥ 1284; 1328; 960; 741 single nuclei for 22y, 75y, 96y and HGPS cell lines respectively across 3 well replicates. Black horizontal bars with asterisks denote statistical significance across donor groups (*** P < 0.001, ** P < 0.01, * P < 0.05; two-sided Mann-Whitney U tests with Bonferroni correction).

Spatial foci analysis revealed distinctly divergent, donor-specific trajectories of chromatin remodeling. Total foci area was greater in both the 96-year-old and HGPS donors, mirroring the diffuse opening seen in replicative aging, whereas the 75-year-old donor exhibited a smaller total foci area but a significantly higher individual foci intensity and net enrichment ratio (Fig. 4C; Fig. S4D-H). This suggests that in addition to widespread structural unraveling, some individuals or cell lineages may experience chronological aging through highly localized increases in chromatin accessibility at specific regulatory hubs, potentially reflecting the activation of age-associated adaptive or stress-response networks (Benayoun et al., 2019; Jänes et al., 2018; Ucar et al., 2017).

Despite these spatial differences, immunofluorescent profiling confirmed a shared underlying erosion of canonical chromatin compartmentalization. We observed an age- and disease-dependent reduction in the constitutive heterochromatin mark H3K9me3 (Fig. 4D; Fig. S4J, K), alongside progressive declines in the active marks H3K4me3 and H3K27ac (Fig. 4F, G; Fig. S4M, N). This concurrent loss of both repressive and activating marks reflects the documented, fundamental degradation of chromatin regulation during aging (Benayoun et al., 2019; Sen et al., 2016). Interestingly, the facultative mark H3K27me3 decreased in fibroblasts from the 75-year-old but accumulated in those from the 96-year-old and HGPS donors, further highlighting phenotypic divergence (Fig. 4E; Fig. S4L). This targeted loss of H3K27me3 tightly correlates with the unique spatial metrics observed in the 75-year-old donor, suggesting that the erosion of facultative heterochromatin may manifest spatially as localized accessible foci in addition to diffuse structural unraveling. Finally, bulk nucleosomal dynamics in these chronological samples diverged from the replicative aging model: total histone H3 decreased with both advanced age and premature aging, while the pan-histone marker tended to increase with age but declined significantly in the HGPS line (Fig. 4H, I; Fig. S4O, P).

Together, these data show that replicative, chronological, and pathological aging are associated with increased global chromatin accessibility and extensive spatial epigenetic remodeling for fibroblasts. The concordance of our ATAC-see metrics with canonical histone mark profiles supports ATAC-see–derived nuclear features as robust, imaging-based biomarkers of cellular aging, providing a powerful tool for image-based screening of geroprotective and disease-modifying interventions.

### ATAC-see captures established multiphasic chromatin remodeling kinetics during OSKM reprogramming

We next tested whether our platform could visualize other chromatin state transitions, such as Yamanaka factor–induced reprogramming and rejuvenation. Because OSKM are potent pioneer transcription factors that physically pry open closed chromatin, we hypothesized that OSKM-driven rejuvenation would induce profound, quantifiable chromatin remodeling detectable by ATAC-see, enabling us to visualize the epigenetic roadmap of reprogramming.

To map the temporal kinetics of this cell fate transition, we examined aged HDFs engineered with doxycycline-inducible OSKM (alongside GFP or wild-type controls) over an 11-day time course. Cells were sampled every one to two days and plated at uniform densities 24 h prior to fixation to eliminate confluence-related artifacts. Multiplexing ATAC-see with immunofluorescence to capture histone modifications, we observed a rapid, continuous increase in global chromatin accessibility between 4 h and 3 days of OSKM induction (Fig. 5A–F and Fig. S5A-F, I, J). This initial burst recapitulated the known pioneer activity of OSKM factors rapidly invading and opening closed chromatin (Fig. 5A) (Soufi et al., 2012). Because early OSKM expression induced a concomitant increase in Hoechst intensity, likely reflecting enhanced dye penetrance into the actively decondensing chromatin, we present both raw (Fig. 5B, C) and DNA-normalized ATAC-see intensities (Fig. S5C, D). Moreover, our multiplexed immunofluorescence confirmed that this early physical opening was driven by an expansion of active chromatin, as we observed an upregulation of the active euchromatin mark H3K4me3 alongside the structural opening of chromatin (Fig. 5B, E and Fig. S5A-D), in stark contrast to the loss of H3K4me3 observed in replicative and chronological aging (Fig. 3H and Fig. 4F). Spatially, average foci area and individual foci intensity expanded dramatically during this early window (Fig. 5D and Fig. S5H), capturing the highly localized, targeted euchromatin expansion expected from pioneer factor binding. In turn, the constitutive heterochromatin mark H3K9me3 was initially variable but remained stable on average (Fig. 5B and Fig. S5D).

**Figure 5.**
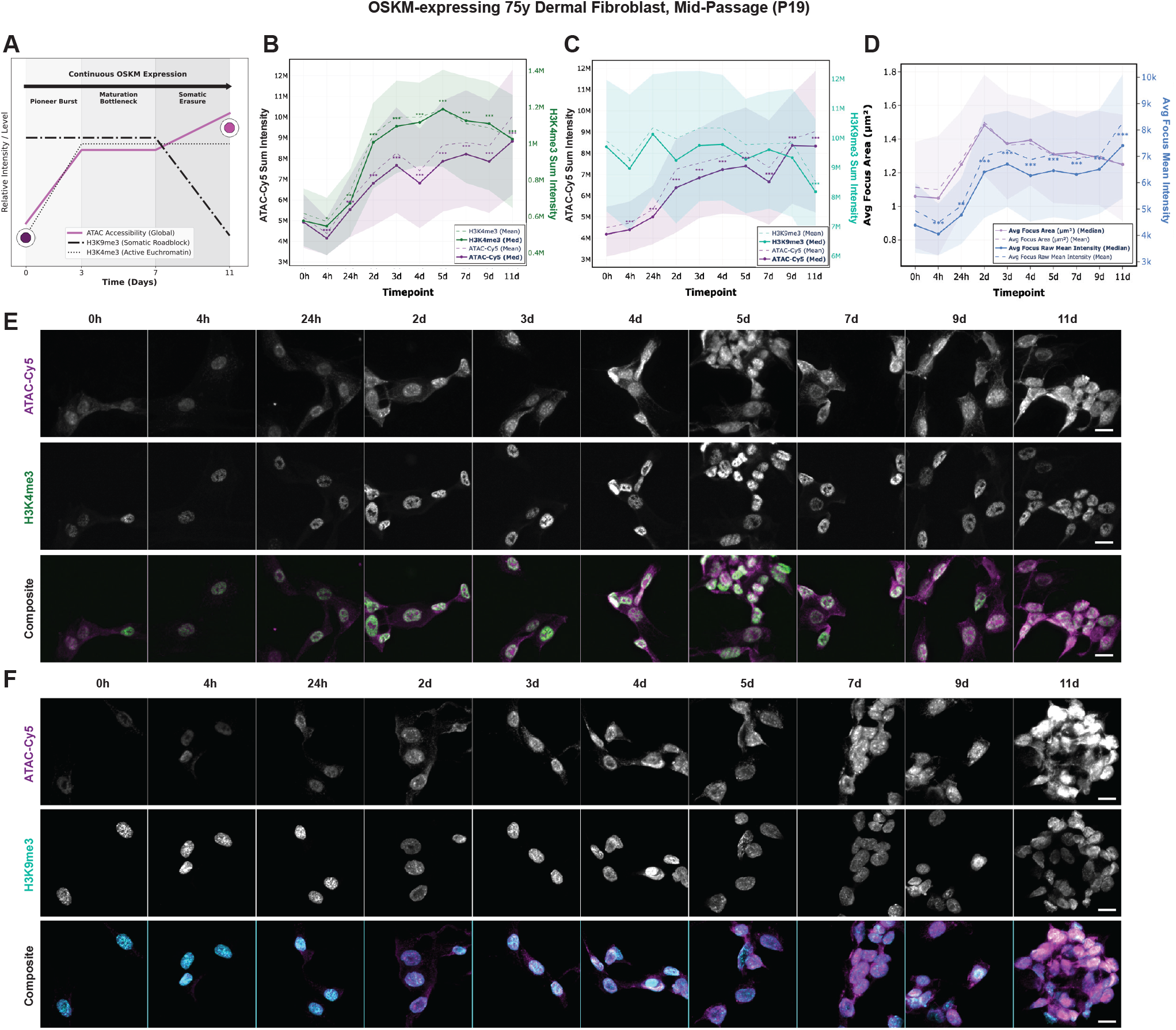
Temporal dynamics of global chromatin accessibility and spatial remodeling during OSKM-induced reprogramming captured by ATAC-see. (A) Schematic model of continuous OSKM expression synthesizing established epigenetic trajectories (Apostolou and Hochedlinger, 2013). The model outlines three biophysical phases: a ‘Pioneer Burst’, a ‘Maturation Bottleneck’, and ‘Somatic Erasure’. The theoretical trajectories of global ATAC accessibility, somatic roadblocks (H3K9me3), and active euchromatin (H3K4me3) are overlaid. (B–D) Quantitative single-nuclei time-course tracking of OSKM-expressing 75-year-old dermal fibroblasts at mid-passage (P19) across an 11-day trajectory. Line graphs display a dual-axis overlay of H3K4me3 (green) and ATAC-Cy5 (purple) Sum Intensities (B), H3K9me3 (cyan) and ATAC-Cy5 (purple) Sum Intensities (B), and subnuclear spatial metrics including Average Focus Area (light purple) and Average Focus Raw Mean Intensity (blue) (D). For all line graphs, solid and dashed lines represent the median and mean values, respectively, while the shaded regions represent the interquartile ranges. (B, C) Data are representative of a single 96-well plate experiment. Each time point represents a pool of 3 replicate wells, with a minimum of n ≥ 706 single nuclei analyzed per condition for H3K4me3 (B) and n ≥ 727 single nuclei for H3K9me3 (C); n ≥ 1553 single nuclei analyzed for foci (D) across 6 well replicates. N ≥ 3 biological replicates. (E–F) Representative high-throughput widefield images capturing the reprogramming time-course from 0h to 11d. Panels highlight the parallel spatial transitions of ATAC-Cy5 with H3K4me3 (E) and H3K9me3 (F), alongside merged composite images. Scale bars, 20µm. Statistical significance is denoted by *** P < 0.001, ** P < 0.01, * P < 0.05; two-sided Mann-Whitney U tests with Bonferroni correction).

As reprogramming progressed into the known maturation bottleneck (days 3–7) (Fig. 5A), the rate of structural opening decelerated and stabilized (Buganim et al., 2012; Polo et al., 2012). Concurrently, H3K4me3 levels plateaued, and nuclear expansion resulted in a relative decline in spatial foci density and occupancy (Fig. 5C and Fig. S5G, H). However, consistent with the kinetics of somatic memory erasure (Apostolou and Hochedlinger, 2013; Chen et al., 2013), a distinct secondary phase emerged between days 7 and 11. An acceleration in chromatin opening and increased individual foci intensity coincided with a substantial reduction in H3K9me3, tracking with the plateau and subsequent decline of sustained OCT4 and SOX2 detection (Fig. 5A-F and Fig. S5B-D). Throughout the entire time course, the facultative heterochromatin mark H3K27me3 progressively increased (Fig. S5E, F), mirroring the expected accumulation of Polycomb-mediated repression required to establish pluripotent bivalency (Polo et al., 2012).

Together, these high-resolution spatial kinetics demonstrate that ATAC-see faithfully captures the established, non-linear epigenetic trajectory of OSKM-driven reprogramming. By mapping the localized, transcriptionally active euchromatin burst of early induction through to the delayed collapse of somatic heterochromatin, ATAC-see provides a powerful imaging-based approach for tracking complex cell fate transitions *in situ* without the need for bulk genomic sequencing.

### ATAC-see tracks chromatin state resolution during transient, rejuvenation-targeted reprogramming

We next investigated whether ATAC-see could capture the resolution of chromatin states during a transient reprogramming paradigm. Continuous OSKM expression ultimately drives the erasure of somatic identity (as observed in our late 11-day timepoints); however, short-term, transient OSK(M) expression can successfully reverse age-associated hallmarks while preserving the original somatic cell lineage (Lu et al., 2020; Ocampo et al., 2016). Based on our temporal kinetic map, we hypothesized that a restricted pulse of OSK(M) would induce the early, transcriptionally active euchromatin burst necessary for epigenetic rejuvenation, while a subsequent recovery period would allow the chromatin to stabilize before reaching the irreversible collapse of somatic heterochromatin.

We subjected aged, late-passage HDFs to a 4-day OSK or OSKM induction, followed by a 4-day withdrawal and recovery phase (Fig. 6A). As expected based on our continuous time course, both OSKM and OSK induction triggered a significant increase in chromatin opening (Fig. 6A-C and Fig. S6A, B). However, only OSKM induced dilation of foci area, whereas both conditions increased mean foci intensity (Fig. 6C, D and Fig. S6D, E). This divergence is consistent with the absence of c-MYC in the OSK condition, as c-MYC functions as a potent global amplifier of transcription and euchromatin expansion (Nie et al., 2012), promoting a more pronounced spatial chromatin relaxation.

**Figure 6.**
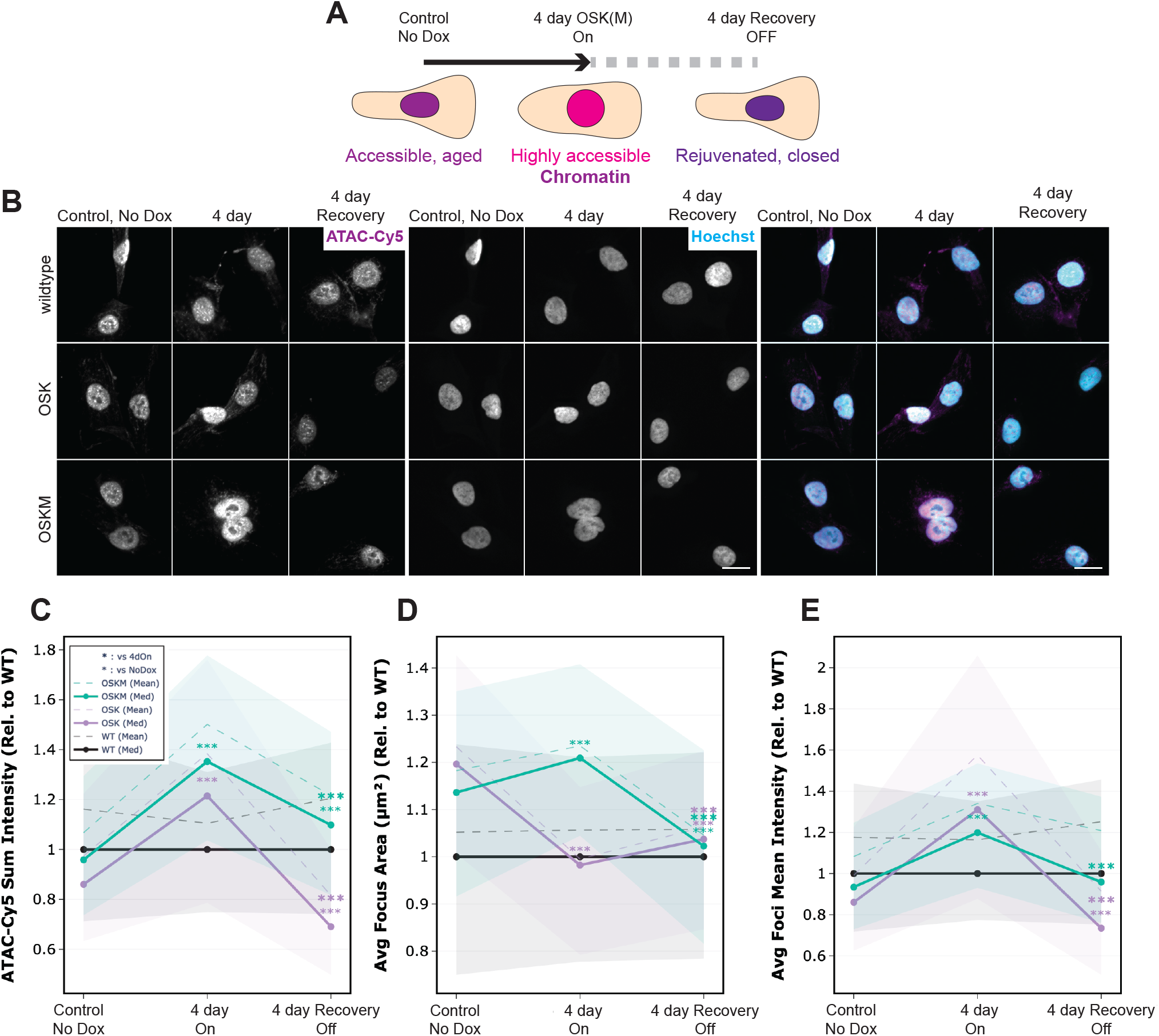
Transient OSKM expression induces reversible global chromatin accessibility and spatial remodeling. (A) Schematic of the transient reprogramming protocol, depicting a 4-day continuous induction ("4 day On") of OSK or OSKM factors followed by a 4-day withdrawal ("4 day Recovery OFF"). The conceptual transition from an aged, accessible state to a highly accessible intermediate, followed by a rejuvenated, closed state, is illustrated. (B) Representative widefield images capturing spatial ATAC-Cy5 dynamics alongside Hoechst counterstaining in wildtype, OSK-, and OSKM-expressing 75-year-old high-passage dermal fibroblasts. Scale bars, 20µm. (C–E) Quantitative tracking of single-nuclei ATAC-see metrics across the transient induction and recovery phases, plotted relative to the matched wildtype (WT) control baseline (value = 1.0, black line), OSK (purple), OSKM (green). Line graphs display Raw Nuclear ATAC Sum Intensity (C), Average Focus Area (D), and Average Focus ATAC Mean Intensity (E). Data are representative of a single 96-well plate experiment, which was performed independently more than 4 times (N = 4 biological replicates). n ≥ 357 single nuclei across 2 well replicates per condition (genotype and time point). Solid and dashed lines represent the median and mean values, respectively, with shaded regions denoting the interquartile ranges. Bold asterisks indicate statistical significance of the treatment condition at a given timepoint relative to the control (No Dox), non-bold asterisks represent significance between 4 day On and Recovery (*** P < 0.001, ** P < 0.01, * P < 0.05; two-sided Mann-Whitney U tests with Bonferroni correction).

Notably, although recovery from both OSK and OSKM expression reduced chromatin accessibility from peak levels, OSK reduced accessibility below the aged baseline (Fig. 6A-D). This suggests that the chromatin landscape partially reverted toward a more youthful, condensed state. Consistent with this, both OSK and OSKM showed reduced foci area (at both the individual and cumulative nuclear levels) and decreased occupancy relative to baseline when normalized to the wild-type controls (Fig. 6D, E and Fig. S6C-L). We further confirmed that these spatiotemporal changes were specific to the Yamanaka factors and not an artifact of transcriptional induction using GFP-inducible controls (Fig. S6M-R).

Together, these findings demonstrate that ATAC-see sensitively captures Yamanaka factor-driven chromatin remodeling in aged cells, providing a robust and highly scalable imaging platform to monitor the extent, trajectory, and reversibility of chromatin opening during partial cellular reprogramming.

### Optimized *in situ* ATAC-see reveals divergent, tissue-specific chromatin aging trajectories

While *in vitro* models provide valuable mechanistic insights, capturing the true physiological complexity of aging requires evaluating chromatin architecture within its native tissue microenvironment. However, adapting transposase-based imaging assays for *in situ* applications presents significant technical hurdles, primarily due to uneven enzyme penetrance and substantial tissue autofluorescence (Deng et al., 2022), and existing *in vitro* ATAC-see protocols failed to yield robust signals in our tissue samples. Drawing on recent advances in spatial ATAC-seq (Deng et al., 2022), we engineered an efficient and robust in situ ATAC-see pipeline optimized specifically for fresh-frozen murine tissue sections (Fig. 7A).

**Figure 7.**
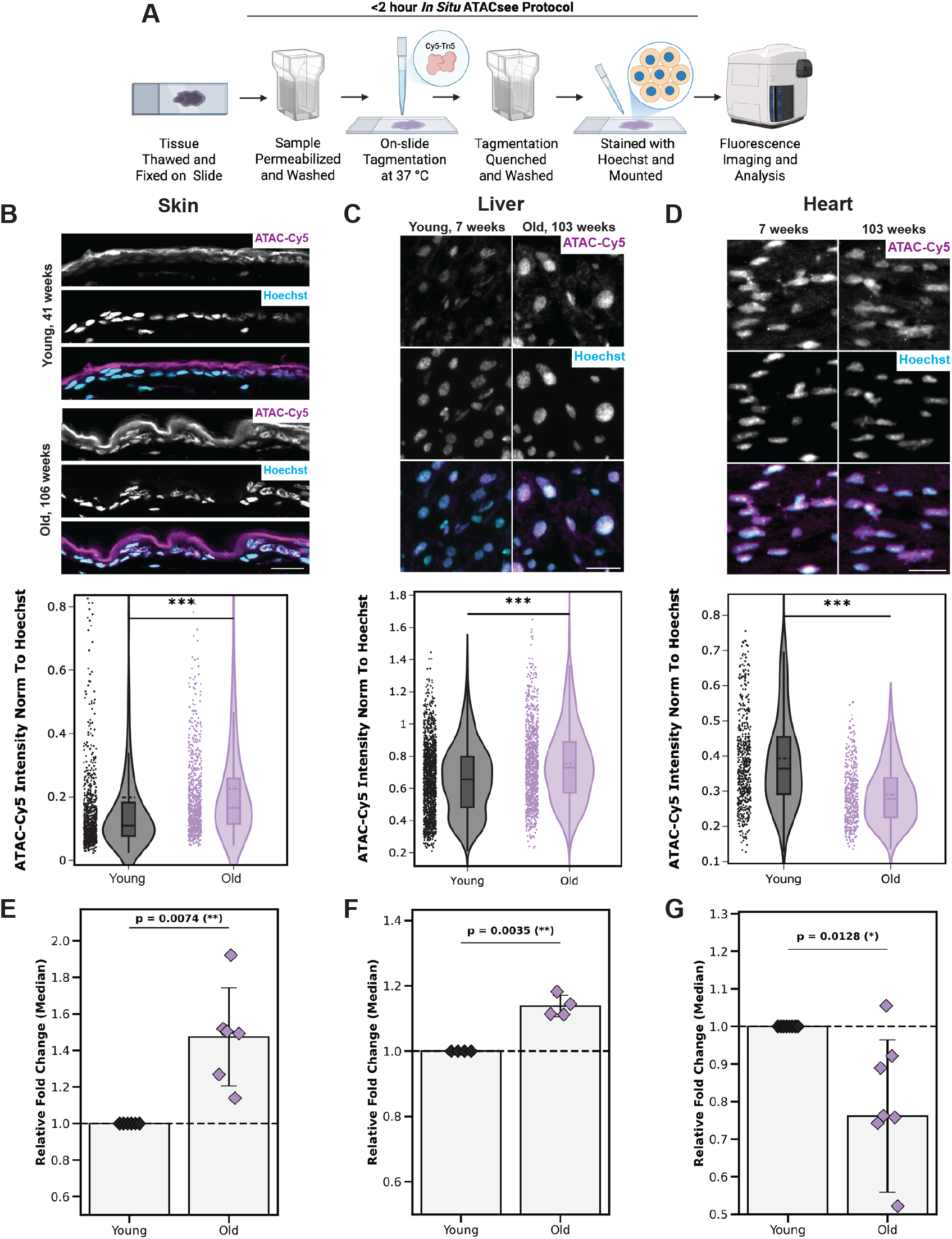
*In situ* ATAC-see maps age-dependent global chromatin remodeling across diverse mammalian tissues. (A) Schematic overview of the optimized, rapid (< 2 hour) *in situ* ATAC-see protocol adapted for cryosectioned mammalian tissue slides. (B–D) Representative confocal images of ATAC-Cy5 (magenta), Hoechst (cyan), and merged channels are displayed alongside corresponding violin plots for single-nuclear intensities of global chromatin accessibility (ATAC-Cy5 Sum Intensity normalized to Hoechst) comparing young and aged tissue sections. Skin (B; 41 weeks vs. 106 weeks), Liver (C; 7 weeks vs. 103 weeks) and Heart (D; 7 weeks vs. 103 weeks). Scale bars, 20µm. Asterisks denote statistical significance across the single-nuclear populations (*** P < 0.001; two-sided Mann-Whitney U tests). (E–G) Relative fold change of the median ATAC-Cy5 intensity norm to hoechst for old versus young samples. Error bars represent s.d., *p* values denoted on graphs determined via independent one-sample t-tests against a non-changing mean of 1.0 (dashed line). Biological replicates, *N* = 6 (Skin), 4 (Liver) and 7 (Heart).

To overcome the unique structural barriers of 3D tissue, we implemented a series of targeted technical modifications. We first stabilized native tissue architecture using a 1% PFA fixation prior to transposase exposure. To achieve sufficient nuclear permeabilization, we applied a 15-min agitated wash in 0.5% Igepal-630 and 0.01% digitonin. Subsequent washes in 0.1% Tween containing 1% bovine serum albumin (BSA) were used to remove excess Igepal. We further optimized the tagmentation buffer by adding 100 mM NaCl and omitting DPBS (Deng et al., 2022) which, together with 1% BSA significantly reduced non-specific background signal and prevented loss of the Tn5 enzyme. Finally, following 60 min of tagmentation at 37°C in a humidified chamber, we employed a stringent 5-min quenching step with 40 mM EDTA, 300 mM NaCl, and 0.005% SDS to remove unintegrated fluorescent oligonucleotides prior to DNA counterstaining and imaging. We applied this modified ATAC-see protocol to a panel of fresh-frozen murine tissues and achieved uniform Tn5 penetrance, preservation of complex cytoarchitecture and minimal background autofluorescence, thereby enabling high-contrast, single-cell visualization of accessible chromatin networks *in situ* (Fig. 7B-D and Fig. S7A, C, E).

To evaluate age-associated chromatin dynamics within tissues, we obtained a small cohort of paired young and aged mouse organs (skin, liver, and heart) and quantified DNA-normalized ATAC-see intensities, accounting for tissue depth-dependent signal loss as well as age-related differences in cell cycle and polyploidy (Donne et al., 2020; Øvrebø and Edgar, 2018). Consistent with our *in vitro* replicative and chronological aging models, we observed a significant global increase in chromatin accessibility in aged skin (106 weeks versus 41 weeks) (Fig. 7B, E and Fig. S7B, E). This progressive epigenetic opening was similarly conserved in aged liver (103 weeks versus 7 weeks) after accounting for the higher proportion of cycling cells in young livers (Fig. 7C, F and Fig. S7C, D). Heart tissue exhibited the variability expected during aging, but showed in contrast, a potential decline in chromatin accessibility, which may align with recent transcriptomic and structural characterizations of the aging heart (Derks and Bergmann, 2020; Gilsbach et al., 2014; Hu et al., 2024; Kirkland et al., 2022) (Fig. 7D and Fig. S7I-L).

These results demonstrate that our newly developed *in situ* ATAC-see protocol enables high-quality imaging of chromatin architecture in complex fresh-frozen sections, and may reveal distinct, organ-specific trajectories of epigenetic aging. This *in situ* capability provides a powerful, highly scalable platform for mapping chromatin accessibility landscapes across the lifespan and evaluating the tissue-specific impacts of systemic rejuvenation therapies.

## Discussion

Structural and biochemical reorganization of the epigenome is a primary driver of cellular dysfunction and a central target for disease and aging interventions. However, a scalable, imaging-based platform capable of resolving these physical chromatin dynamics at single-cell resolution within native spatial contexts is lacking. Genomic assays like ATAC-seq provide invaluable base-pair resolution but sacrifice spatial architecture, while traditional immunofluorescence captures individual histone modifications but cannot directly quantify the physical accessibility of the DNA itself. In this study, we comprehensively optimize and extend the ATAC-see protocol to establish a standardized imaging platform for quantifying chromatin accessibility in fixed cells and fresh-frozen tissues. By systematically calibrating fixation and tagmentation kinetics *in vitro,* we addressed the key technical limitations that have historically hindered the broader application of *in situ* transposase assays. We demonstrate that this platform captures the spatiotemporal chromatin trajectories of chronological, pathological, and replicative aging, dynamically tracks Yamanaka factor-driven chromatin remodeling, and successfully maps chromatin accessibility within intact mammalian tissues. Ultimately, this optimized ATAC-see platform establishes spatial chromatin accessibility as a practical, quantitative imaging biomarker candidate of aging and rejuvenation, while preserving full compatibility with downstream genomic sequencing.

Our optimized pipeline addresses a methodological gap by balancing spatial resolution with high-throughput scalability. While recent super-resolution adaptations like 3D ATAC-PALM provide unprecedented, nanometer-scale mapping of chromatin networks in cultured cells (Xie et al., 2020), the requirement for highly specialized instrumentation and complex sample preparation limits their utility for larger-scale studies. Conversely, recent efforts to adapt ATAC-see for high-throughput flow cytometry (Ishii et al., 2024) offer rapid single-cell accessibility readouts but fundamentally sacrifice sub-nuclear spatial context. Furthermore, as our systematic parameter exploration in fixed fibroblasts revealed, protocols relying on short tagmentation windows fail to overcome the diffusion barrier of even lightly crosslinked nuclei, resulting in reduced signal-to-noise ratios. By systematically calibrating a 60-min tagmentation window with 1% PFA fixation, our platform captures quantitative spatial metrics, such as foci size, intensity, and net enrichment, that are not possible with traditional flow cytometry, while remaining compatible with standard automated microscopy and sequencing.

This spatial sensitivity enabled us to characterize the "passive" chromatin unraveling of aging and the "active" regulatory remodeling of rejuvenation in our cell models. During replicative aging and in primary fibroblasts from aged and Hutchinson-Gilford progeria syndrome (HGPS) donors, age-associated opening manifests largely as a diffuse, uncoordinated unraveling. Rather than a targeted regulatory expansion, this diffuse opening reflects a disordered loss of higher-order chromatin architecture and the progressive degradation of heterochromatin boundaries (Cecco et al., 2013; Villeponteau, 1997). This structural entropy disrupts nuclear compartmentalization, resulting in a loss of spatial contrast and a "flattened" epigenetic landscape that is simultaneously depleted of canonical active marks (like H3K4me3) and repressive marks (like H3K9me3) (Benayoun et al., 2019; Sen et al., 2016).

We also demonstrate that the ability to multiplex optimized ATAC-see with targeted immunofluorescence provides a unique lens for uncovering potential biophysical mechanisms driving epigenetic divergence. For instance, our spatial analysis revealed that chronological aging may also give rise to localized accessible regions. By directly correlating this spatial topography with a loss of H3K27me3, our imaging platform is consistent with the hypothesis that the age-associated erosion of Polycomb-mediated repression can cause poised, bivalent regulatory hubs to erroneously open (Dozmorov, 2015; Hahn et al., 2014; Signal et al., 2024). Ultimately, this demonstrates that ATAC-see, when paired with canonical histone profiling, can directly relate specific biochemical epigenetic failures to the physical reorganization of the aging nucleus. Conversely, we observed that reprogramming via OSKM induction triggers an active, highly regulated expansion of euchromatin. ATAC-see visualized this phenomenon as a pioneer factor-driven burst of accessible foci, synchronized with a significant upregulation of H3K4me3.

Mapping the continuous kinetics of OSKM induction allowed us to visualize the established epigenetic roadmap of reprogramming *in situ*, delineating the early pioneer burst from the delayed reduction of somatic heterochromatin (H3K9me3). This temporal map informed our transient reprogramming paradigm, revealing that upon the withdrawal of OSK after four days, the chromatin landscape not only resolves from its hyper-accessible peak but partially condenses below the aged baseline. This structural stabilization into a more tightly regulated, youthful state, achieved without crossing the critical threshold of somatic memory erasure (Apostolou and Hochedlinger, 2013; Ocampo et al., 2016), provides further evidence supporting transient OSK as a rejuvenating intervention and ATAC-see as a potential platform for evaluating further interventions. Furthermore, the ability to rapidly quantify drug-induced chromatin remodeling following TSA treatment, positions optimized ATAC-see as a highly sensitive, image-based metric for compounds targeting epigenetic erosion.

Building on our robust *in vitro* framework, we successfully extended optimized ATAC-see to map chromatin states across diverse *in vivo* microenvironments. While spatial ATAC-seq (Deng et al., 2022) has advanced tissue epigenomics, we found that substituting fluorescent adapters into sequencing-based permeabilization protocols was inadequate for imaging. These conditions degraded nuclear integrity and resulted in high mitochondrial background and inter-tissue variability. Consequently, our *in situ* pipeline, incorporating morphological stabilization, optimized permeabilization, and targeted signal quenching, was necessary for high-quality and robust ATAC-see application in tissues. Applying this, we confirmed that epigenetic aging is not a uniform phenomenon. While fresh-frozen skin and liver sections exhibited the chromatin opening similar to that observed *in vitro*, the aging heart showed a potential reduction in chromatin accessibility. This divergence may be consistent with recent work detailing the unique biology of the aging myocardium, including H1.0 upregulation (Hu et al., 2024), limited accessibility at key transcriptional program sites (Kirkland et al., 2022), and increased structural silencing associated with polyploidization (Derks and Bergmann, 2020; Gilsbach et al., 2014). Furthermore, our observations align with the emerging consensus from large-scale epigenomic atlases. For example, a recent landmark single-cell ATAC-seq atlas encompassing 21 mammalian organs revealed that while aging drives organism-wide epigenomic remodeling, these changes follow highly tissue-specific, cell-type-specific, and sex-dependent regulatory trajectories (Lu et al., 2026). Parallel findings in model organisms, such as the Aging Fly Cell Atlas in *Drosophila*, have similarly demonstrated that different tissues age at fundamentally different rates with distinct transcriptomic and epigenetic signatures (Lu et al., 2023). Indeed, comparative genomic studies have shown that while some organs, like the murine liver, undergo a massive age-related expansion in promoter accessibility, other tissues, such as the immune and hematopoietic compartments, undergo widespread chromatin compaction (Bozukova et al., 2022). Our *in situ* ATAC-see protocol adds a spatial and physical dimension to complement these sequencing-based observations.

### Limitations and Future Directions

Despite its spatial capabilities, ATAC-see is an optical assay and does not yield the sequence-specific regulatory information provided by ATAC-seq. Therefore, this platform is not intended to replace genomic sequencing, but rather to complement it as a first-line imaging assay. Future studies may leverage this platform to map the spatial distribution of rejuvenated cells within whole organs, potentially accelerating the translation of epigenetic interventions from the bench to the clinic.

## Methods

### Cell culture and interventions

#### Primary human dermal fibroblasts (HDFs)

GM05294, GM08401, GM00731 and AG11498 were ordered from Coriell Institute. Cells were cultured in DMEM + 15% FBS + 1X NEAA (non-essential amino acids; Cat# 11140050) and maintained at 37°C and 5% CO_2_ in a humidified incubator. All cell lines were routinely tested for mycoplasma contamination.

#### Long term cell culture and *in vitro* aging model

For the BJ fibroblast *in vitro* aging model, cells were maintained in continuous culture and serially passaged upon reaching 70–80% confluence to prevent contact inhibition. At each passage, cells were washed with PBS and detached using TrypLE, counted using an automated cell counter, and re-seeded at a constant density of 6,600 cells/cm², equivalent to 500,000 cells per T75 flask. The aging trajectory was tracked chronologically, and representative cohorts of cells were cryopreserved at regular intervals to establish a longitudinal bank. For downstream assays, “Low passage” baseline cells were defined as those from the early expansion phase (P6–P10). “Mid passage” cells were defined as P20–P22 (approximately 4–6 weeks post-expansion), while “High passage” cells were defined as pre-senescent at P26–P28 (approximately 6 weeks of further culture). High passage cells remained proliferative, albeit at a notably reduced rate characteristic of late-stage replicative exhaustion.

#### Lentiviral transduction

Coriell GM08401 cells were transduced with a polycistronic OSKM, OSK or GFP cassette under the control of a Tet-On (TRE) promoter, alongside an rtTA transactivator and puromycin selection at low passage using a lentiviral delivery system.

Lentiviral particles were generated using the TransIT-VirusGEN® LV Transfection Kit (Mirus Bio) in combination with Lenti-X™ Concentrator (Takara Bio). Briefly, packaging cells were seeded in 15 cm dishes at an approximate density of 1.5×10^7^ cells per dish in 30 mL of complete growth medium and incubated overnight to reach 80–95% confluence. For each 15 cm dish, a total of 30 µg of plasmid DNA, comprising 15 µg of the transgene transfer plasmid and 15 µg of an optimized packaging plasmid premix, was diluted into 3 mL of VirusGEN® LV Complex Formation Solution. TransIT-VirusGEN® Transfection Reagent (90 µL) was added to the DNA mixture, vortexed gently, and incubated at room temperature for 15 to 60 min to allow complex formation. The complexes were added dropwise to the cultures. Between 18 and 24 hours post-transfection, 3 mL of VirusGEN® LV Enhancer was added directly to each dish. Viral supernatants were harvested 48 hours post-transfection and clarified by centrifugation at 500 × *g* for 10 min to remove cellular debris. Clarified supernatant was mixed with Lenti-X™ Concentrator at a 3:1 volumetric ratio and incubated overnight at 4°C. The mixture was centrifuged at 1,500 × *g* for 45 min at 4°C to pellet the virions. After a secondary brief spin to completely aspirate residual media, the off-white viral pellet was gently resuspended in 360 µL of sterile PBS, divided into single-use aliquots, and flash-frozen at −80°C.

To empirically determine optimal transduction efficiencies and minimize viral toxicity, functional titration was performed prior to bulk cell line engineering. Target cells were seeded in 24-well plates at empirically optimized densities (up to 90% confluence) to promote survival during infection. Serial dilutions of the concentrated lentivirus (ranging from 0.04 to 25 µL/mL) were prepared in culture media supplemented with TransPlus™ Virus Transduction Enhancer (Alstem) at a concentration of 2 µL/mL. Cells were exposed to the virus-enhancer mixture, and transduction efficacy versus cell viability was monitored with puromycin at 1ug/ml. 25% survival was used for optimal selection.

#### Doxycycline-inducible reprogramming

To initiate continuous or transient reprogramming, stable cell lines (GM08401, encoding OSKM, OSK, GFP, or wild-type controls) were seeded in 6-well plates (120,000 cells per well). Expression was induced by supplementing the culture media with 1 µM Doxycycline (Dox). To maintain stable induction, Dox-containing media was refreshed every 48 hours. Cells were passaged as required following the onset of induction to prevent contact inhibition and over-confluence. 24 hours prior to terminal fixation, cells from all conditions were detached using TrypLE, counted, and re-seeded into 96-well optical imaging plates (CellVis, glass bottom) at a precise density of 5,000 cells per well in their respective control or Dox-containing media. For acute early-induction time point, 4 hours, the control media was replaced with Dox media at the respective time point prior to fixation. Cells were then washed once with HBSS (containing CaCl^2^ and MgCl^2^) (Cat# 14025134), then fixed as per the respective ATACsee or Immunofluorescence protocol.

#### Pharmacological treatments

Cells were seeded in 96-well plates 24 hours prior to treatment to allow for adherence and to ensure equal cell density. Trichostatin A (TSA) stock solutions in DMSO were diluted in culture media to final concentrations ranging from 62.5 nM to 1 µM and applied to the cells immediately following the aspiration of the standard growth media. Control cells were treated with an equivalent volume of DMSO in media. To prevent solvent-induced cytotoxicity, the final concentration of DMSO in all treated and control wells was maintained strictly below 0.01% (v/v). Following a 4-hour incubation with either TSA or vehicle control, the media was removed, and cells were washed with PBS (containing CaCl₂ and MgCl₂) prior to fixation.

### Animal models and tissue processing

#### Animal models and tissue acquisition

All animal procedures and tissue harvests were conducted in accordance with the NIH Guide for the Care and Use of Laboratory Animals and approved by the Institutional Animal Care and Use Committee (IACUC) at Altos Labs.

Tissues were harvested from strictly male C57BL/6J mice at predetermined end-of-life time points. Prior to tissue collection, all mice were group-housed (maintained in groups of at least two animals per cage to prevent isolation stress) under standard specific pathogen-free (SPF) conditions, with *ad libitum* access to standard chow and water.

To evaluate age-associated spatial chromatin dynamics, tissues were collected from distinct cross-sectional age cohorts. The liver was harvested from young, 7 week, and aged, 103 week, male mice, n = 4 pairs. Young heart samples were harvested from two cohorts of male mice that were 7-12 weeks and 103-104 weeks, n = 7 pairs. Skin samples were acquired from a separate cohort of mature (41 weeks) and aged (106 weeks) male mice, n = 6. With the exception of heart tissue, upon euthanasia, tissues were immediately embedded in Optimal Cutting Temperature (OCT) compound (Tissue-Tek). Crucially, to control for downstream processing variance, young and aged organs of the same tissue type were paired and co-embedded within the same OCT block. The blocks were then snap-frozen in a 2-methylbutane (isopentane) bath cooled with dry ice and stored at −80°C. For one heart sample cohort, liquid-nitrogen snap frozen hearts were embedded in OCT post storage at −80°C. Hearts had been independently embedded but were prepared on the same slides as the other organs.

#### Cryosectioning

Prior to sectioning, OCT-embedded fresh-frozen tissue blocks were equilibrated to the cryostat chamber temperature (20°C). Tissues were sectioned at a thickness of 12 µm or 16 µm for Heart and Liver or 18µm for Skin, using a cryostat. Because most young and aged tissues were co-embedded, they were simultaneously sectioned and captured onto the same Superfrost Plus charged microscope slides to ensure identical thickness, adherence, and spatial processing conditions. A subset of heart tissues were sectioned from independent blocks and arranged on the same slide during sectioning. Following sectioning, the slides were briefly air-dried to promote tissue adherence and immediately stored at −80°C until required for ATAC-see staining and imaging.

### Tn5 Transposase production and assembly

#### Tn5 hyperactive double mutant w/ c-terminal mxe intein and chitin binding domain

MITSALHRAADWAKSVFSSAALGDPRRTARLVNVAAQLAKYSGKSITISSEGS**K**AMQEGAYRFIRNPNV SAEAIRKAGAMQTVKLAQEFPELLAIEDTTSLSYRHQVAEELGKLGSIQDKSRGWWVHSVLLLEATTFRTVGLLHQEWWMRPDDPADADEKESGKWLAAAATSRLRMGSMMSNVIAVCDREADIHAYLQDKLAHNERFVVRSKHPRKDVESGLYLYDHLKNQPELGGYQISIPQKGVVDKRGKRKNRPARKASLSLRSGRITLKQGNITLNAVLAEEINPPKGETPLKWLLLTSEPVESLAQALRVIDIYTHRWRIEEFHKAWKTGAGAERQR MEEPDNLERMVSILSFVAVRLLQLRESFT**P**PQALRAQGLLKEAEHVESQSAETVLTPDECQLLGYLDKG KRKRKEKAGSLQWAYMAIARLGGFMDSKRTGIASWGALWEGWEALQSKLDGFLAAKDLMAQGIKICITGDALVALPEGESVRIADIVPGARPNSDNAIDLKVLDRHGNPVLADRLFHSGEHPVYTVRTVEGLRVTGTANHPLLCLVDVAGVPTLLWKLIDEIKPGDYAVIQRSAFSVDCAGFARGKPEFAPTTYTVGVPGLVRFLEAHHRDPDAQAIADELTDGRFYYAKVASVTDAGVQPVYSLRVDTADHAFITNGFVSHATGLTGLNSGLTTNPGVSAWQVNTAYTAGQLVTYNGKTYKCLQPHTSLAGWEPSNVPALWQLQ

#### Tn5 hyperactive double mutant with n-terminal protein-a/g and c-terminal mxe intein and chitin binding domain

MSLKDDPSQSANLLSEAKKLNESQAPKADNKFNKEQQNAFYEILHLPNLNEEQRNGFIQSLKDDPSQSANLLAEAKKLNDAQAPKADNKFNKEQQNAFYEILHLPNLTEEQRNGFIQSLKDDPSVSKEILAEAKKLNDAQAPKTTYKLVINGKTLKGETTTEAVDAETAERHFKQYANDNGVDGEWTYDDATKTFTVTEKPEVIDAS ELTPAVDDDKEFGGGGSGGGGSGGGGSGGGGSHMITSALHRAADWAKSVFSSAALGDPRRTARLVNVAAQLAKYSGKSITISSEGSKAMQEGAYRFIRNPNVSAEAIRKAGAMQTVKLAQEFPELLAIEDTTSLSYRHQVAEELGKLGSIQDKSRGWWVHSVLLLEATTFRTVGLLHQEWWMRPDDPADADEKESGKWLAAAATSRLRMGSMMSNVIAVCDREADIHAYLQDKLAHNERFVVRSKHPRKDVESGLYLYDHLKNQPELGGYQISIPQKGVVDKRGKRKNRPARKASLSLRSGRITLKQGNITLNAVLAEEINPPKGETPLKWLLLTSEPVESLAQALRVIDIYTHRWRIEEFHKAWKTGAGAERQRMEEPDNLERMVSILSFVAVRLLQLRESFTPPQALRAQGLLKEAEHVESQSAETVLTPDECQLLGYLDKGKRKRKEKAGSLQWAYMAIARLGGFMDSKRTGIASWGALWEGWEALQSKLDGFLAAKDLMAQGIKICITGDALVALPEGESVRIADIVPGARPNSDNAIDLKVLDRHGNPVLADRLFHSGEHPVYTVRTVEGLRVTGTANHPLLCLVDVAGVPTLLWKLIDEIKPGDYAVIQRSAFSVDCAGFARGKPEFAPTTYTVGVPGLVRFLEAHHRDPDAQAIADELTDGRFYYAKVASVTDAGVQPVYSLRVDTADHAFITNGFVSHATGLTGLNSGLTTNPGVSAWQVNTAYTAGQLVTYNGKTYKCLQPHTSLAGWEPSNVPALWQLQ

#### Tn5 expression

The genes for hyperactive Tn5 double mutant (Tn5) and the pAG-tagged Tn5 double mutant (pAG-Tn5) were synthesized and cloned into separate empty pET3a vectors using NdeI and BamHI restriction sites. Plasmid was transformed following a standard protocol into BL21DE3 E. coli (Cat# EC0114) and grown at 37C overnight on LB agar plates containing 100ug/ml Ampicillin (Teknova Cat# L1002). An 80mL LB starter culture containing 100ug/ml carbenicillin (Corning Cat# 46-100-RG) in a 250ml baffled flask was inoculated with a single isolated colony and grown at 30C overnight. The next morning, 10mL of dense starter culture was sub-cultured into 1L of LB containing 100ug/ml carbenicillin in a 2.7L baffled flask and grown at 37C until an OD between 0.6-0.8 was reached, upon which protein induction was initiated by the addition of 1mM IPTG (Fisher CAT# J17886.14) and the incubator temperature dropped to 22C for 18h. Cells were then harvested by centrifugation at 8,000g using a JLA 8.1000 rotor in a Beckman centrifuge and cell pellets were stored at −80C until purification day.

#### Tn5 E. coli lysis and purification

The morning of the lysis, cell pellets were thawed on ice for approximately 2h then resuspended in 10ml per gram of HEGX buffer (50mM HEPES pH 7.2, 800mM NaCl, 0.8mM EDTA, 10% v/v glycerol, 0.2% Triton X-100) containing protease inhibitor cocktail (cat# p8849). After complete resuspension, the cells were disrupted by sonication on ice (5 sec on / 5 sec off cycle, 60% amplitude, 10 min total sonication). Lysate was then transferred to centrifuge tubes and spun at 40,000g for 20 min at 4C using a JA 25.50 rotor in an Avanti centrifuge. The supernatant was then transferred to a beaker and bulk DNA was precipitated by the dropwise addition of 10% v/v polyethyleneamine 1,200 (PEI) to a final concentration of 0.1% v/v or until the solution became cloudy. The solution was then transferred back to centrifuge tubes and spun at 40,000g for 20 min at 4C using a JA 25.50 rotor. The clarified supernatant was then decanted off the DNA pellet and was ready for the purification.

The following purification was carried out completely at 4C, with all buffers pre-chilled. The clarified supernatant from the previous step was poured onto a gravity-flow column containing 0.5ml of HEGX-equilibrated chitin resin (cat# S6651L) per gram of cell pellet lysed. The lysate was allowed to flow through the column twice and collected to ensure maximal binding. After binding, the resin was washed by 30 column volumes (CV) of HEGX buffer, then the intein elution was initiated by the addition of 2 CV of elution buffer (HEGX containing 100 mM dithiothreitol [DTT; freshly dissolved before addition]). 1 CV of the elution buffer was allowed to flow through the column, then the column flow was ceased by stopcock and 1 CV of buffer was left on the column for 72 h to ensure intein cleavage went to completion and liberated the Tn5. After 72 h, the remaining elution buffer was collected, and the column was washed with 1 CV of HEGX to remove any residual liberated Tn5. The protein was concentrated to 10 mL and injected onto a HEGX-equilibrated 320 mL HiLoad S200 size exclusion column attached to an Akta Pure FPLC system. The column was washed with 1CV of HEGX and all fractions were collected by an automated fractionator. The Tn5 peak eluted at roughly 0.6-0.8 CV and fractions corresponding to the entire peak were collected and pooled. Protein concentration was determined by BCA assay and concentrated to a final concentration of 1.5mg/ml (roughly 27uM). The protein was then dialyzed overnight into storage buffer (50 mM HEPES, 800 mM NaCl, 0.2 mM EDTA, 20% v/v glycerol, 2 mM DTT), aliquoted into eppendorf tubes (256 uL each), and flash frozen in liquid nitrogen before being stored at −80C until transposome loading.

#### Mosaic end sequences and transposome loading

Tn5ME-Reverse (R): /5Phos/CT GTC TCT TAT ACA CAT CT

Tn5ME-A-Cy5 (A): /5Cy5/TC GTC GGC AGC GTC AGA TGT GTA TAA GAG ACA G

Tn5ME-B-Cy5 (B): /5Cy5/GT CTC GTG GGC TCG GAG ATG TGT ATA AGA GAC AG

The 5’ Cy5 can be replaced with Cy3 or Alexafluor488 fluorophores, but Cy5 was generally preferred.

Single-stranded DNA oligonucleotides were ordered from IDT containing the sequences above with the respective fluorophores on forward strands (reverse strand unlabelled) and dissolved to stock concentrations of 200 uM with nuclease-free H2O. The following protocol is an example of how to make 1,028 uL of 20x (3.4uM) of fluorescently loaded transposomes for ATACsee.

The double-stranded A-R (Tn5ME-A-Cy5 hybridized with Tn5ME-Reverse) oligo was created by combining 12.8 uL of 200 uM Tn5ME-A-Cy5 (A) with 12.8 uL of 200uM Tn5ME-Reverse (R) and diluting with 25.6 uL of annealing buffer (10 mM Tris-HCl, pH 8.0) to a volume of 51.2 uL and concentration of 50uM total DNA in a PCR tube. The double-stranded B-R oligo was created by combining 12.8 uL of 200 uM Tn5ME-B-Cy5 with 12.8 uL of 200 uM Tn5ME-Reverse and diluting with 25.6 uL of annealing buffer (10 mM Tris-HCl, pH 8.0) to a final concentration of 50 uM total DNA in a PCR tube. The A-R and B-R oligonucleotides were respectively annealed into double-stranded DNA using a BioRad T100 thermocycler with the following settings: First the oligonucleotides were heated to 95 C for 5 min, then cooled down to 65 C at 0.1 C/second and held there for 5 min, followed by another cooldown to 4 C at 0.1 C/second where it was held until the next step.

The separate double-stranded oligonucleotides (A-R and B-R) were then diluted to 128uL by adding 76.8 uL of 85% v/v glycerol in H2O to a concentration of 20 uM DNA, then both A-R and B-R solutions were transferred into a 1.5 mL Eppendorf tube (128uL of A/R and 128ul of B/R; total volume of 256 uL). The combined A-R and B-R oligonucleotides were further diluted with 640 uL of 50% glycerol to a volume of 886 uL. −80C frozen Tn5 (27 uM stock) was thawed on ice then 128ul of the stock was added to the mixture, mixed thoroughly by pipetting, and allowed to load with the DNA for 1 h at room temperature in the dark (final volume of 1028 uL). The final glycerol concentration for the transposomes was 47% v/v glycerol and were stored at −30C until use on the day of ATACsee experiments.

### Optimized ATAC-see protocols

#### *In vitro* ATAC-see (fixed cells)

Cells cultured in 96-well glass-bottom plates (Cell Vis) were washed once with HBSS containing calcium and magnesium (HBSS + Ions; Thermo Fisher, Cat# 14025134) at room temperature. Cells were subsequently fixed using freshly diluted 1% (w/v) paraformaldehyde (PFA; Cat# 50-980-494) in HBSS + ions for 10 min at room temperature. Any alternative fixation conditions are explicitly detailed in the respective figure legends. Fixation was quenched via three sequential washes with HBSS + ions. Following fixation, plates were immediately processed for ATAC-see, subjected to immunofluorescent staining, or stored at 4°C for up to one week.

Transposomes (3.4 µM stock) were diluted 20-fold to a final working concentration of 170 nM in 1× Tagment DNA (TD) buffer (10 mM TAPS, pH 8.0, 5 mM MgCl₂, 10% *N,N*-dimethylformamide; prepared from a 2× stock) supplemented with 0.1% IGEPAL CA-630, and mixed thoroughly by pipetting. The working transposome solution was sonicated in a bath sonicator at 37 kHz for 60 seconds at 80% amplitude. To remove residual aggregates, the solution was passed through a 0.22 µm syringe filter (Cat# SLGVR33RS).

A combined single-step cell permeabilization and tagmentation (PermNTag) was initiated by adding 100 µL of the filtered working transposome solution to each well of the fixed cells. Plates were incubated at 37°C for the designated duration (typically 1 hour). To control for background fluorescence and determine specificity, 20mM EDTA was added to the tagmentation solution for control samples. Tagmentation was quenched by washing the cells twice with calcium/magnesium-free HBSS (HBSS w/o Ions; Cat# 14065056) supplemented with 40 mM EDTA and 0.01% SDS. Cells were then washed three times with standard HBSS at room temperature, counterstained with Hoechst 33342 (1:1000 dilution, Invitrogen) for 15–25 min, washed three additional times with HBSS, and stored at 4°C until microscopic imaging.

For experiments combining ATAC-see with immunofluorescence (IF), the complete IF protocol was performed prior to the ATAC-see PermNTag step. This sequence prevents the SDS used in the tagmentation quenching buffer from adversely affecting epitope antigenicity or causing antibody degradation.

#### *In situ* ATAC-see (fresh-frozen tissues)

Slides contained two spatially separated regions for young and old tissues to capture inter-organ variability and to enable internal normalization respectively. Slides stored at −80°C were first warmed at room temperature for 2 min. Tissues were fixed on the slide by immersing in 1% formaldehyde in PBS (containing ions for increasing adhesion) for 5 min. Following fixation, the solution was removed, and the slides were washed three times with PBS (with ions). To facilitate enzyme penetration, tissues were permeabilized with PBS supplemented with 0.5% Igepal and 0.01% digitonin for 15 min at room temperature, with agitation on an orbi-shaker (30 rpm). The slides were subsequently washed twice for 5 min each on an orbi-shaker using a wash buffer consisting of PBS with 0.1% Tween-20 and 1% BSA.

During washes, the transposition reaction mix was prepared containing 1× TD buffer, 100 mM NaCl, 0.1% Tween-20, 0.01% digitonin, 1% BSA, and 1X Cy5-conjugated Tn5 transposase (from a 20X stock, final concentration of 170nM), brought to volume with purified water. Care should be taken to ensure that the tissues are sufficiently immersed allowing for small amounts of evaporation. The slides were then incubated at 37 °C for 60 min inside a humidified chamber with gentle agitation (20-30 rpm) to allow for robust chromatin tagmentation. We utilized black western Biorad boxes, but any vessel for sufficient coverage can be used that also protects against light and evaporation. To control for background fluorescence and determine specificity, 20mM EDTA was added to the tagmentation solution for control samples.

To immediately halt the transposition reaction, slides were transferred to a histology slide holder and quenched for 5 min using a stop buffer of 0.005% SDS, 300 mM NaCl, and 40 mM EDTA in PBS. Following quenching, the solution was removed, and the slides were washed three times with PBS to remove detergent. Nuclei were counterstained directly on the slide using Hoechst (1:1,000 dilution in PBS) for 15-25 min. The slides were washed three final times with PBS in the holder and subsequently mounted using DAKO mounting medium. The mounted tissues were left to cure overnight at room temperature in the dark.

Special note: To account for batch-to-batch variation observed during tissue processing, likely stemming from non-specific Tn5 absorption by plastic or OCT, variable vessel coverage, or evaporation, an internal control (young tissue) was maintained on every slide. Furthermore, differences observed between distinct organs processed in the same batch may arise from true biological variance or tissue-dependent differences in permeabilization efficiency. Therefore, permeabilization parameters may be further optimized on an organ-specific basis.

### Immunofluorescence

#### Standard immunofluorescence

Following 4% fixation for 10 min, samples were rinsed 3X with HBSS, then permeabilized using 0.5% Triton X-100 in HBSS for 15 min at room temperature. Permeabilization was followed by three 10-min washes in HBSS containing 0.1% Triton X-100. To minimize non-specific antibody binding, samples were blocked for 1 hour at room temperature using Antibody Dilution buffer (AbDil; 150 mM NaCl, 20 mM Tris-HCl [pH 7.4], 0.1% Triton X-100, 2% bovine serum albumin, and 0.1% sodium azide in H₂O, sterile filtered).

Samples were subsequently incubated overnight at 4 °C with primary antibodies diluted in AbDil. The following unconjugated primary antibodies were utilized: rabbit anti-H3K27me3 (Sigma-Aldrich, Cat# 07-449; 1:200), rabbit anti-H3K9me3 (Active Motif, Cat# 61013; 1:400), mouse anti-H3K4me3 (Active Motif, Cat# 61379; 1:500), rabbit anti-H3K27ac (Abcam, Cat# ab4729; 1:1,000), Histone H3 (Abcam-1791, 1:1000) and Histone H1-4 (MAB-3422, 1:1000). For transcription factor profiling, directly conjugated primary antibodies targeting Sox2 (Sox2-488; Invitrogen Cat#53-9811-82, 1:100) and Oct4 (Oct3/4-647; BD Cat#560307, 1:100).

Following primary incubation, the plates were washed three times for 10 min each in HBSS with 0.1% Triton X-100. For unconjugated targets, samples were then incubated with corresponding species-specific secondary antibodies (diluted 1:200-400 in AbDil from central stocks that were prepared per manufacturers instructions, then diluted 1:1 with glycerol) and Hoechst diluted 1:1000, for 1.5 hours at room temperature.

To ensure a clean imaging background, the plates underwent three 10-min washes in HBSS with 0.1% Triton X-100, followed by three final washes in standard HBSS to completely remove all detergents prior to imaging.

#### Multiplexed-immunofluorescence

IF was performed first prior to ATAC-see on 1% PFA, 10 min, fixed samples. Triton-X-100 was substituted for Igepal-NP40. Following 3X HBSS washes, cells were permeabilized with 0.5% Igepal-NP40 in HBSS for 15 min, then further washed with 0.1% Igepal-HBSS for 5 min, 3 times. Cells were blocked with HBSS containing 2% BSA and 0.1% Igepal for 1 hour. Cells were then incubated with the respective primary antibodies (concentrations noted above) for 1 hour at room temperature. Cells were then washed 3 x 10 min with 0.1% Igepal-HBSS. The secondary antibody was prepared and incubated as above for 1 hour. Subsequently, cells were washed

#### Staining protocol

Detail how IF was performed *after* ATAC-see quenching. Include blocking buffers, primary antibody incubation times/temps, and secondary antibodies with 0.1% Igepal-HBSS for 5 min, 3 times before proceeding to the ATAC see tagmentation protocol.

### High-throughput imaging

#### Image acquisition

Automated widefield epifluorescence imaging was performed utilizing a Nikon Eclipse Ti2 inverted microscope integrated into a BioPipeline high-throughput imaging system. Prepared 96-well plates were automatically loaded and scanned using a Nikon CFI Plan Apochromat Lambda D 20X air objective (NA 0.80) paired with a Photometrix Kinetix 25mm camera, Lumencor Spectra III light engine, Semrock Brightline individual or seven-filter “Pinkel” penta-band filter cubes, and a Cairn OptoSpin filter wheel with individual 32mm Semrock Brightline emission filters. To ensure a robust dynamic range for the quantitative assessment of both intense localized foci and diffuse nucleoplasmic signals, all images were captured at a 16-bit pixel depth. Multi-channel acquisition was fully automated via NIS-Elements software. To compensate for micro-fluctuations in plate topography during automated scanning, Nikon’s Perfect Focus System (PFS) was employed to establish a stable baseline focal plane. Images were acquired at multiple spatially distributed fields of view per well. At each field of view, images were captured as Z-stacks consisting of 9 optical slices with a 0.8 µm step size.

High-resolution fluorescence imaging of fresh-frozen murine tissue sections was performed manually using a Zeiss LSM 900 laser scanning confocal microscope. Fields of view were acquired using a Plan Apochromat 20X objective (NA 0.8) with an additional 2x optical zoom, yielding a frame size of 1024 x 1024 pixels. To ensure a high dynamic range for quantitative intensity measurements, all images were captured at a 16-bit pixel depth. Z-stacks were acquired for each field of view using a 1 µm step size. 3 images were obtained per section, totaling 3 technical replicates each. Total Z-stack depths ranged from 18 to 22 µm, adjusted per region to accommodate local variations in tissue section topography and thickness.

### ATAC-Seq after imaging

To demonstrate the compatibility of ATAC-see with downstream sequencing, the ATAC-see protocol was first performed on cells seeded in 12-well glass-bottom plates (Cellvis), alongside matched EDTA-treated negative controls (20 mM EDTA). Following imaging on a Nikon Eclipse Ti2 inverted microscope as described above, HBSS was aspirated and replaced with a reverse-crosslinking solution adapted from (Chen et al., 2016), consisting of 50 mM Tris-HCl, 1 mM EDTA, 1% SDS, 0.2 M NaCl and 250 ug/ml proteinase K. Cells were physically detached using a cell scraper, transferred to microcentrifuge tubes, and incubated overnight at 65°C with agitation at 1,000 rpm in a thermomixer. DNA was subsequently purified using the MinElute Reaction Cleanup Kit (Qiagen, Cat#: 28204) according to the manufacturer’s instructions. Eluted DNA was amplified using 11, 14, and 17 PCR cycles with NEBNext® HiFi 2× PCR Master Mix (NEB, M0541S) and pre-mixed Illumina DNA/RNA UD Indexes (Set A; cat. no. 20091646), following the PCR protocol described in (Buenrostro et al., 2013). Library quality was assessed by Agilent TapeStation, and products from all three cycle numbers were pooled and purified using 1.8× SPRI beads (Beckman Coulter, cat. no. B23318). Sequencing was performed on an Illumina NextSeq 2000 (P3 flow cell) with paired-end 50 + 50 bp reads and 10 bp dual-indexing (50/10/10/50 cycle configuration).

For downstream analysis, sequencing data were processed using the nf-core/cutandrun pipeline (v2.1.2) with default parameters, except that MACS2 peak calling was run in narrow peak mode on reads pooled across all six replicates.

### Automated image analysis and quantification

#### Image pre-processing and segmentation

To establish a standardized 2D focal plane from the acquired multi-slice Z-stacks, raw images were processed into single-plane projections utilizing the Extended Depth of Focus (EDF) algorithm within Nikon NIS-Elements software for *in vitro* studies. For *in situ* studies, Sum or single slices were generated using FIJI. Subsequent image analysis and single-cell feature extraction were performed using Arivis Pro 4.2.0 and 4.3.0 software. Single-nucleus segmentation was achieved by applying the integrated Cellpose deep-learning tool (utilizing the pre-trained cyto3 model) directly to the Hoechst counterstain channel. The resulting verified nuclear masks were then applied across all corresponding experimental channels to extract bulk morphological and fluorescence intensity metrics.

#### Automated spatial foci detection

Subnuclear spatial chromatin architecture was quantified using the Blob Finder tool within Arivis. To specifically preserve and enhance high-intensity localized signals against diffuse nucleoplasmic background, images were first pre-processed using a Top-Hat filter. To ensure optimal detection fidelity across biologically distinct populations, the expected blob radius was empirically optimized and visually verified for each specific dataset. Using a defined parent-child relationship parameter, all detected foci (blobs) were spatially assigned to their respective individual parent nuclei, enabling the extraction of foci count, area, and mean intensity strictly within the nuclear boundary.

#### Data normalization and statistical analysis

Extracted single-cell metrics were exported as CSV files for downstream quantitative analysis. To account for technical and biological variations, such as focal depth, cell cycle phase, and ploidy, in cases specified in text, raw sum intensities for ATAC-Cy5 and immunofluorescence marks were normalized to total DNA content on a per-nucleus basis, utilizing the respective Hoechst sum intensity. Data analysis, normalization, and statistical evaluation were performed utilizing custom scripts in Jupyter Notebooks (Python version 3.11.4). Array operations and data structuring were executed using the numpy and pandas libraries, while data visualization was generated utilizing plotly, seaborn and matplotlib. All statistical analyses were performed using the scipy.stats module. Statistical significance for pairwise comparisons between independent in vitro experimental groups was determined using the non-parametric two-sided Mann-Whitney U test. To account for multiple comparisons across extensive combinatorial matrices (e.g., dose-response gradients), p-values were explicitly adjusted using the Bonferroni correction. For variance analysis across more than two independent groups (e.g., time-course progression), the Kruskal-Wallis H-test was employed, followed by appropriate pairwise post-hoc testing utilizing the scikit-posthocs library. *p* values were determined as “*” for < 0.05, “**” for < 0.01 and “***” for < 0.001. For in vitro analysis, 2-3 wells were imaged per condition in most cases, and the experiment was repeated across several biological replicates for confidence. Statistical power analyses were not performed *a priori* to determine sample sizes. As segmentation and intensity extraction were executed using these fully automated computational pipelines, user blinding during image acquisition and analysis was not required.

### Use of Artificial Intelligence tools

The large language model Gemini (Google) was used to adapt code for data visualization and to refine manuscript text. AI was not used for data analysis or scientific interpretation. The authors have reviewed and verified all AI-assisted code, text, and references for accuracy and take full responsibility for the content of this manuscript.

## Acknowledgements

We would like to thank Eric Griffis and Judy Martin of the Imaging hub, and Tatyana Makushok, Maria Burdyniuk, Valentina Pedoia from the Biovision Hub for supporting microscopy and image analysis training. We also thank Christa Caneda and Garret Bishop of the Cell Modeling and Engineering Hub for supporting all in vitro operations and Angela Denn of the Histology Hub. We thank Nasun Ha and the genomics hub for supporting genomic sequencing. Finally, we thank the operations and facilities teams for enabling all daily activities. We thank Francesco Della Valle for comments on the manuscript.

## Author Contributions

N.K. and M.A.C. conceived the project, performed the core experiments, and wrote the original manuscript. N.K. and Y.Y analyzed and interpreted the quantitative data. M.A.C. designed and conducted the biochemical protocols for Tn5 expression and purification. M.J. and V.L. generated and provided the transient OSK- and OSKM-expressing cells utilized for the ATAC-see experiments. B.S., Y.Y. and F.M. performed the harvesting and sectioning of the murine tissues. P.M.-C. critically reviewed and edited the manuscript. P.M.-C., J.C.I.B. and Z.L. supervised the research program.

## Data and Resource availability

The raw and processed sequencing data generated in this study have been deposited in the NCBI Gene Expression Omnibus (GEO) under accession number GSE338065. Due to proprietary institutional restrictions, raw imaging datasets are not publicly deposited but are available from Altos Labs, Inc. upon reasonable request for non-commercial research purposes, subject to a standard institutional Data Sharing Agreement (DSA).

Custom image analysis workflows were executed using commercially available software (Nikon NIS-Elements, Arivis Vision4D) as described in the Methods. Custom Python scripts and Jupyter Notebooks utilized for downstream data normalization, spatial metrics calculation (including the Foci Net Enrichment ratio), statistical analysis, and data visualization will be made available upon reasonable request for non-commercial research purposes, subject to a standard institutional Data Sharing Agreement (DSA).

