## Supplemental Figures for "Visualizing the Epigenetic Landscape of Aging and Cellular Reprogramming: Optimized ATAC-see for Cells and Tissues"

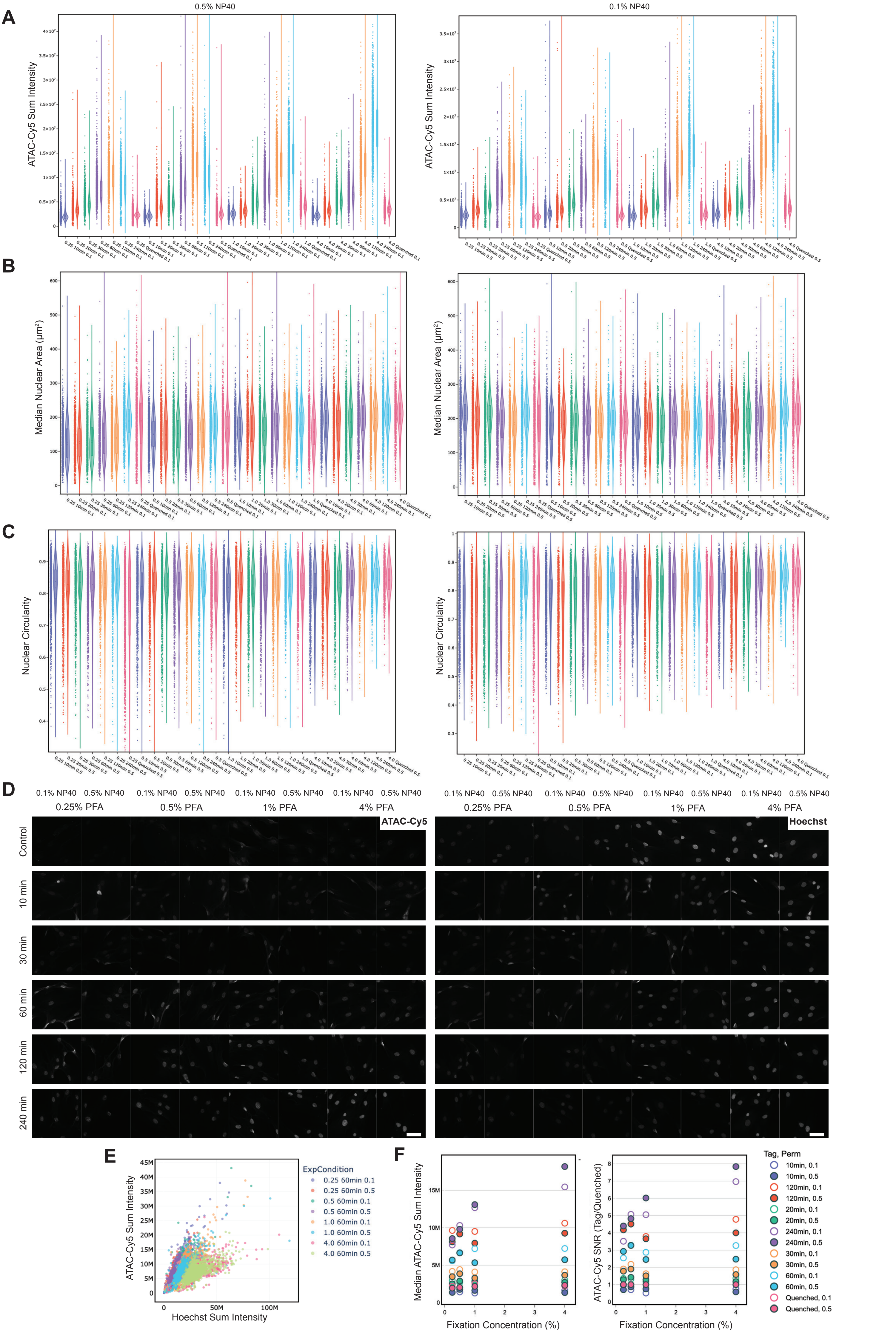
A
0.5% NP40
0.1% NP40
ATAC-Cy5 Sum Intensity
B
Median Nuclear Area (µm²)
C
Nuclear Circularity
D
0.1% NP40
0.5% NP40
0.25% PFA
0.5% PFA
1% PFA
4% PFA
ATAC-Cy5
Hoechst
Control
10 min
30 min
60 min
120 min
240 min
E
ATAC-Cy5 Sum Intensity
Hoechst Sum Intensity
ExpCondition
0.25 60min 0.1
0.25 60min 0.5
0.5 60min 0.1
0.5 60min 0.5
1.0 60min 0.1
1.0 60min 0.5
4.0 60min 0.1
4.0 60min 0.5
F
Median ATAC-Cy5 Sum Intensity
ATAC-Cy5 SNR (Tag/Quenched)
Fixation Concentration (%)
Tag, Perm
10min, 0.1
10min, 0.5
120min, 0.1
120min, 0.5
20min, 0.1
20min, 0.5
240min, 0.1
240min, 0.5
30min, 0.1
30min, 0.5
60min, 0.1
60min, 0.5
Quenched, 0.1
Quenched, 0.5

**Figure S1 | High-resolution single-cell distributions of ATAC-see optimization metrics.** (A–C) Single-cell distribution profiles (violin plots) detailing ATAC-Cy5 sum intensity (A), median nuclear area (B), and nuclear circularity (C) across the complete combinatorial matrix of PFA fixation (0.25%, 0.5%, 1%, 4%), permeabilization (0.1%, left or 0.5% NP40/Igepal, right), and tagmentation times (10 to 240 min, plus quenched controls) for data presented in Fig. 1. (D) Extended panels of representative uncropped fields of view demonstrating the ATAC-Cy5 signal-to-noise ratio and nuclear preservation across the indicated fixation, permeabilization, and tagmentation conditions. Scale bars, 60 µm. (E) Scatter plot showing positive correlation of ATAC-Cy5 and Hoechst (DNA) sum intensities across conditions, highlighting cell cycle and DNA content influence ATAC-see intensities. (F) Summary scatter plots aggregating median ATAC-Cy5 total intensity and SNR across the fixation gradients with extended, 18-min fixation, and at low cell density, reinforcing the selection of intermediate fixation (1% PFA) and a 60-min tagmentation time for optimal assay performance. Data shown is representative from a single 96-well plate experiment, using a minimum of $n \geq 158$ single nuclei per condition.

**75y, Dermal Fibroblast, GM08401 - Trichostatin A**

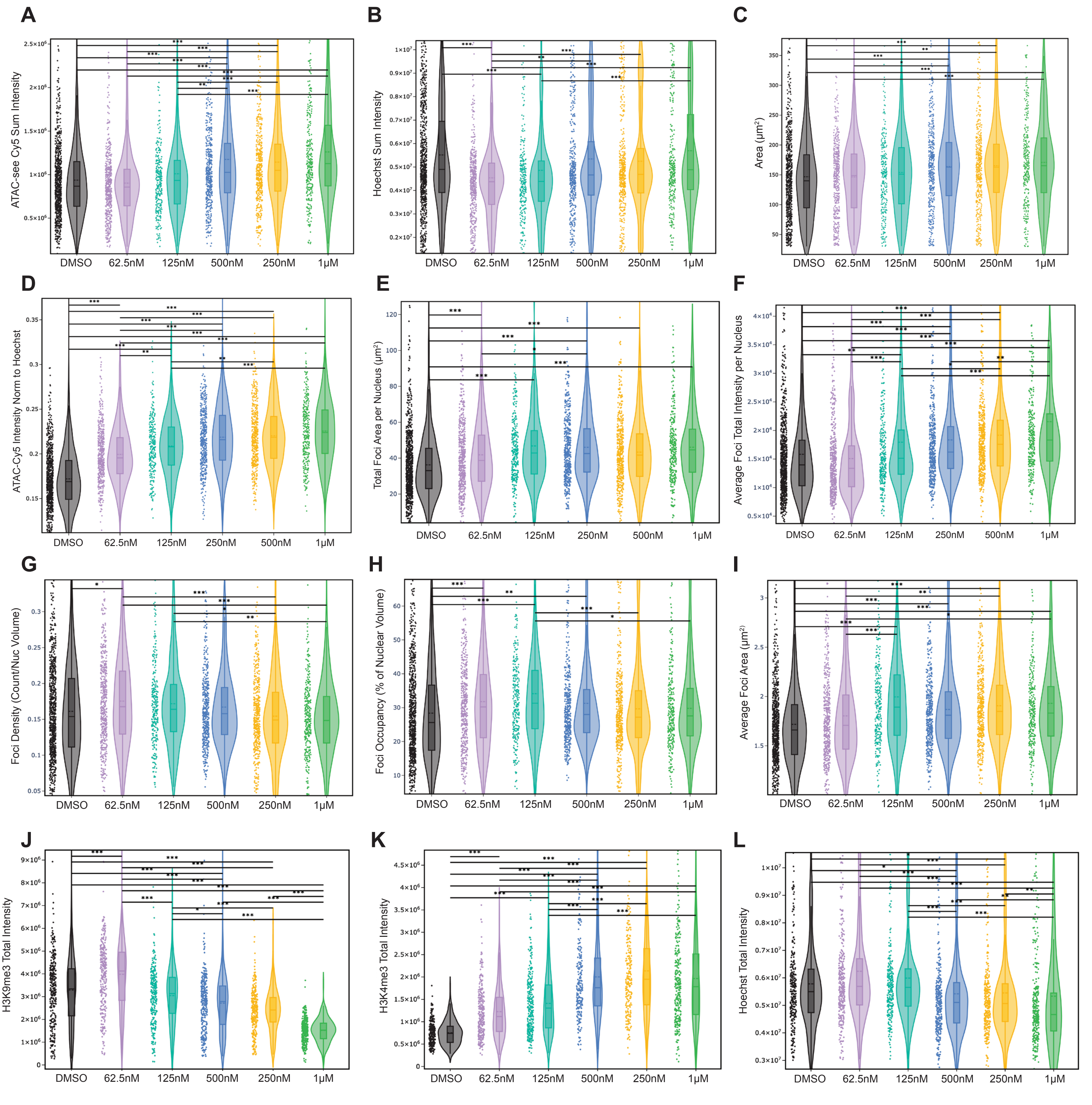

**Figure S2 | Raw intensity distributions and extended spatial metrics during TSA-induced decondensation.** All data represents 75-year-old primary human dermal fibroblasts (GM08401) treated with a TSA dose-response (DMSO, 62.5 nM, 125 nM, 250 nM, 500 nM, 1 μM). (A–B) Single-nuclei distributions of unnormalized, raw ATAC-see Cy5 Total Intensity (A) and corresponding total Hoechst Total Intensity (B) prior to DNA-content normalization. (C) Median nuclear area ($\mu m^2$). (D) DNA-normalized ATAC-Cy5 Intensity. (E-I) Violin plots for subnuclear spatial metrics, including Total Foci Area per nucleus ($\mu m^2$) (E), Average Foci Total intensity per nucleus (F), Foci Density (Count/Nuclear Volume) (G), total Foci Occupancy as a percentage of nuclear volume (H), and Average Foci Area ($\mu m^2$) (I). (J-L) Raw, unnormalized total intensity distributions for parallel immunofluorescence validation of the H3K9me3 (J) and H3K4me3 (K) histone marks and DNA intensity (L). Data are representative of a single 96-well plate experiment, which was performed independently more than 3 times ($N \geq 3$ biological replicates). A minimum of $n \geq 272$ single nuclei for (A-I) and $n \geq 207$ single nuclei for (J-L) were analyzed per condition. Statistical significance across treatment groups: *** $P < 0.001$, ** $P < 0.01$; * $P < 0.05$; two-sided Mann-Whitney U tests with Bonferroni correction.

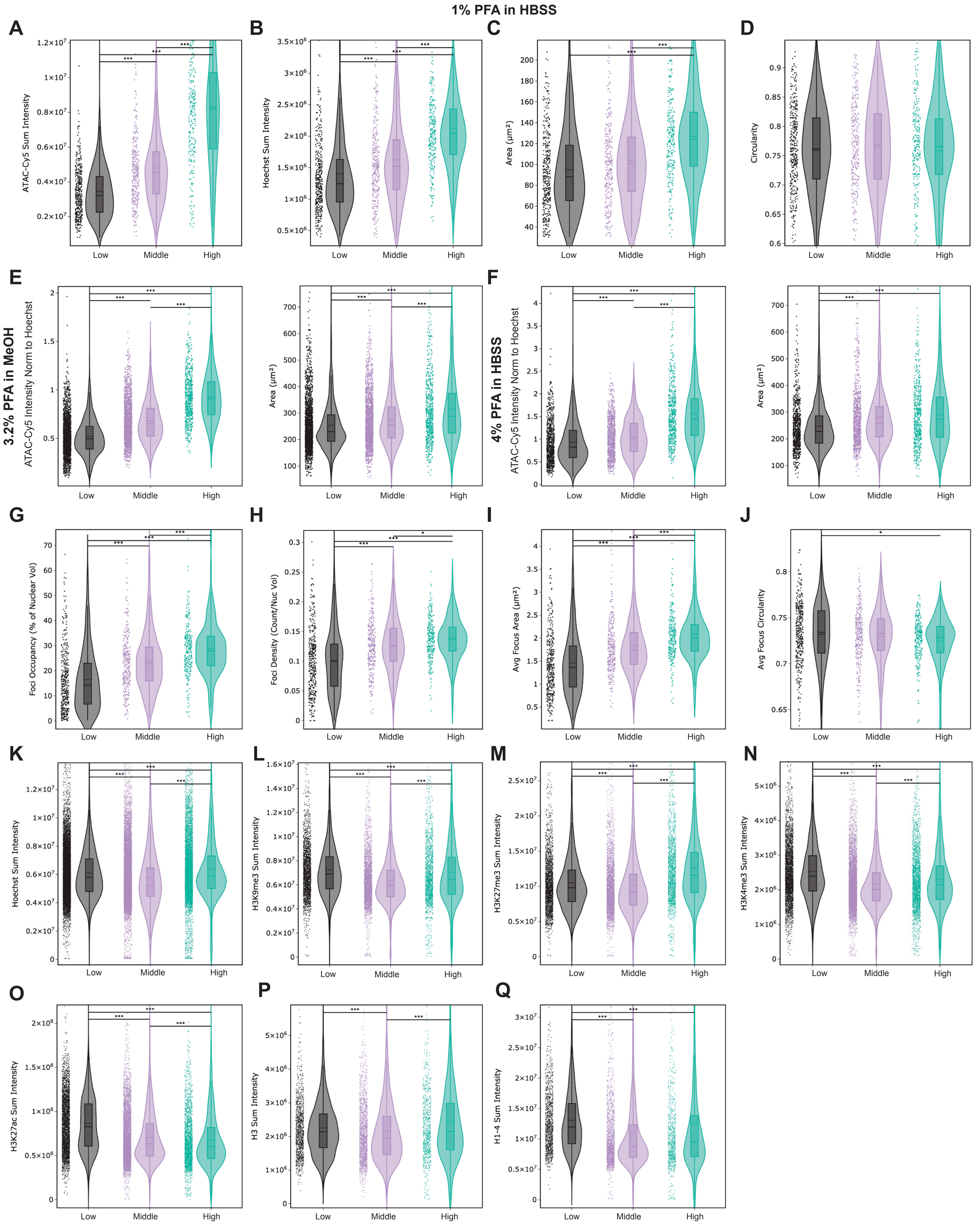
1% PFA in HBSS
A
ATAC-Cy5 Sum Intensity
B
Hoechst Sum Intensity
C
Area (µm²)
D
Circularity
E
3.2% PFA in MeOH
ATAC-Cy5 Intensity Norm to Hoechst
Area (µm²)
F
4% PFA in HBSS
ATAC-Cy5 Intensity Norm to Hoechst
Area (µm²)
G
Foci Occupancy (% of Nuclear Vol)
H
Foci Density (Count/Nuc Vol)
I
Avg Focus Area (µm²)
J
Avg Focus Circularity
K
Hoechst Sum Intensity
L
H3K9me3 Sum Intensity
M
H3K27me3 Sum Intensity
N
H3K4me3 Sum Intensity
O
H3K27ac Sum Intensity
P
H3 Sum Intensity
Q
H1-4 Sum Intensity
Low
Middle
High

**Figure S3 | Unnormalized intensity distributions, morphological shifts, and fixation condition comparisons during continuous *in vitro* aging.** (A–D) Raw, unnormalized single-nuclei metrics for the primary experimental condition (1% PFA in HBSS), including ATAC-Cy5 Sum Intensity (A), Hoechst Sum Intensity (B), Nuclear Area (C), and Nuclear Circularity (D) across Low, Middle, and High passage fibroblasts. (E–F) Validation of passage-induced global chromatin accessibility (ATAC-Cy5 Intensity normalized to Hoechst) utilizing alternative fixation protocols: 3.2% PFA in Methanol ($n$ = 2083; 1562; 522 single nuclei for low, medium and high passage respectively across 3 well replicates) (E) and 4% PFA in HBSS ($n$ = 750; 797; 445 single nuclei for low, medium and high passage respectively across 3 well replicates) (F). (G–H) Extended spatial metrics detailing subnuclear ATAC-Cy5 architecture, including Foci Occupancy as a percentage of nuclear area (G) and Foci Density (H). (I–J) Comparison of Average Focus Area (I) and Average Focus Circularity (J) across passaging. (A-D, G-H) Data are representative of a single 96-well plate experiment, which was performed independently more than 3 times ($N \geq 3$ biological replicates). $n$ = 376; 280; 243 single nuclei for low, medium and high passage respectively across 3 well replicates. (K) Raw Hoechst Sum Intensity distributions corresponding to the parallel immunofluorescence assays. (L–Q) Unnormalized sum intensity distributions for all parallel immunofluorescence targets across the passaging timeline, including H3K9me3 (L), H3K27me3 (M), H3K4me3 (N), H3K27ac (O), Histone H3 (P), and pan-Histone marker, H1-4 (Q). Data are representative of a single 96-well plate experiment, $n \geq$ 1015; 1471; 639 single nuclei for low, medium and high passage respectively across 3 well replicates. Statistical significance: *** $P < 0.001$, ** $P < 0.01$, * $P < 0.05$; two-sided Mann-Whitney U tests with Bonferroni correction.

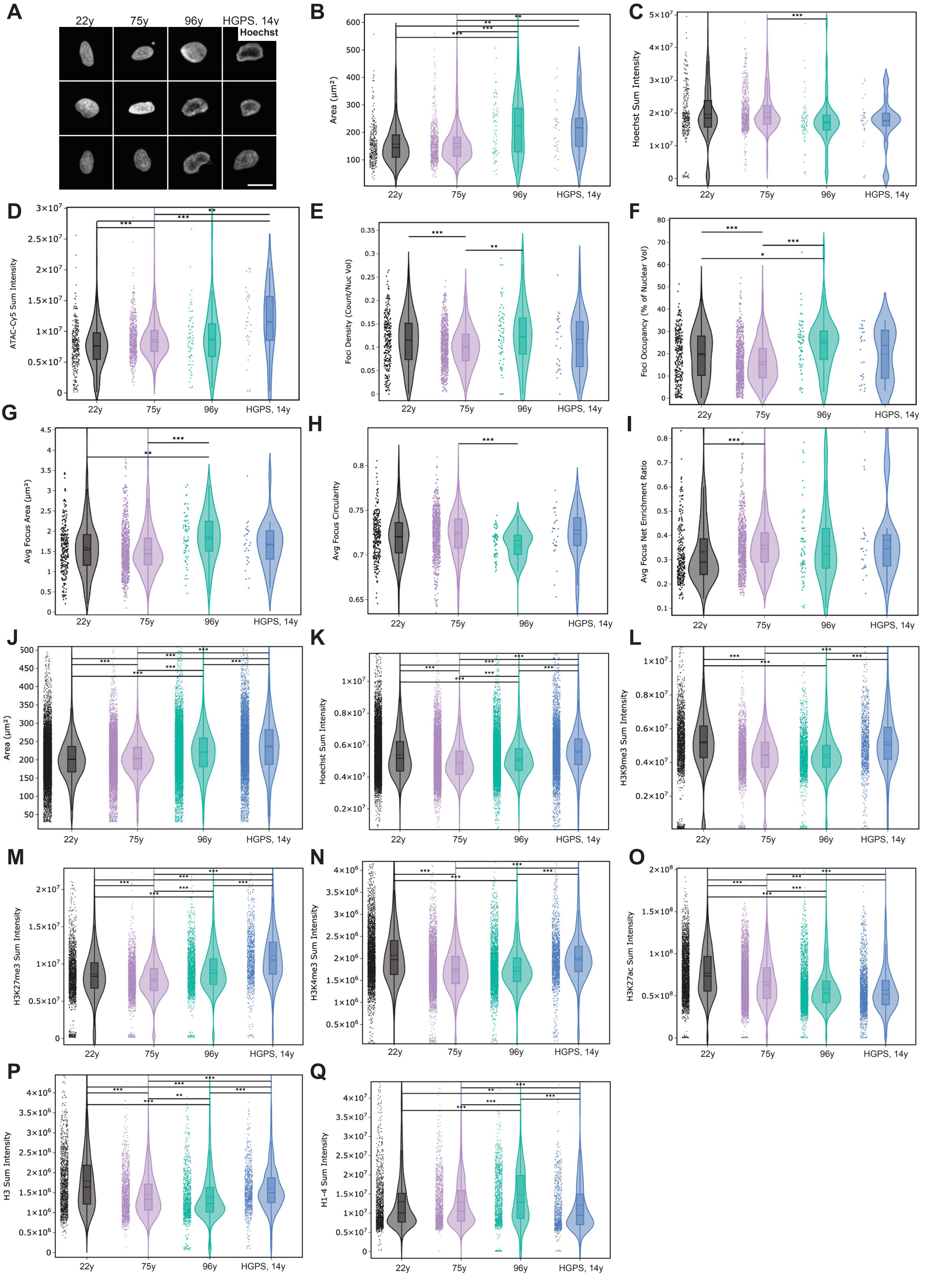
A
22y
75y
96y
HGPS, 14y
Hoechst
B
Area (μm²)
C
Hoechst Sum Intensity
D
ATAC-Cy5 Sum Intensity
E
Foci Density (Count/Nuc Vol)
F
Foci Occupancy (% of Nuclear Vol)
G
Avg Focus Area (μm²)
H
Avg Focus Circularity
I
Avg Focus Net Enrichment Ratio
J
Area (μm²)
K
Hoechst Sum Intensity
L
H3K9me3 Sum Intensity
M
H3K27me3 Sum Intensity
N
H3K4me3 Sum Intensity
O
H3K27ac Sum Intensity
P
H3 Sum Intensity
Q
H1-4 Sum Intensity

**Figure S4 | Unnormalized intensity distributions, morphological shifts, and extended spatial metrics across physiological and premature aging.** (A) Three representative nuclei with Hoechst counterstaining corresponding to the primary donor fibroblast lines: 22y, 75y, 96y, and HGPS (14y). Scale bars, 20μm. (B–C) Single-nuclei distributions of Nuclear Area (B) and total Hoechst Sum Intensity (C) for the ATAC-see experimental plates. (D–H) Extended spatial metrics detailing subnuclear ATAC-Cy5 architecture across the donor cohort, including Foci Density (Count/Nuc Vol) (D), Foci Occupancy as a percentage of nuclear volume (E), Average Focus Area (F), Average Focus Circularity (G), and the Average Focus Net Enrichment Ratio (H). Data are representative of a single 96-well plate experiment, which was performed independently more than 3 times ($N \geq 3$ biological replicates). $n \geq 305$; 704; 75; 36 single nuclei for 22y, 75y, 96y and HGPS cell lines respectively across 2 well replicates. (I–J) Raw Nuclear Area (I) and Hoechst Sum Intensity (J) distributions corresponding to the parallel immunofluorescence assay plates. (K–P) Unnormalized sum intensity distributions for all parallel immunofluorescence targets, including H3K9me3 (K), H3K27me3 (L), H3K4me3 (M), H3K27ac (N), Histone H3 (O), and pan-Histone H1-4 (P). Data are representative of a single 96-well plate experiment, $n \geq 1284$; 1328; 960; 741 single nuclei for 22y, 75y, 96y and HGPS cell lines respectively across 3 well replicates. Statistical significance is denoted by *** $P < 0.001$, ** $P < 0.01$, * $P < 0.05$; two-sided Mann-Whitney U tests with Bonferroni correction).

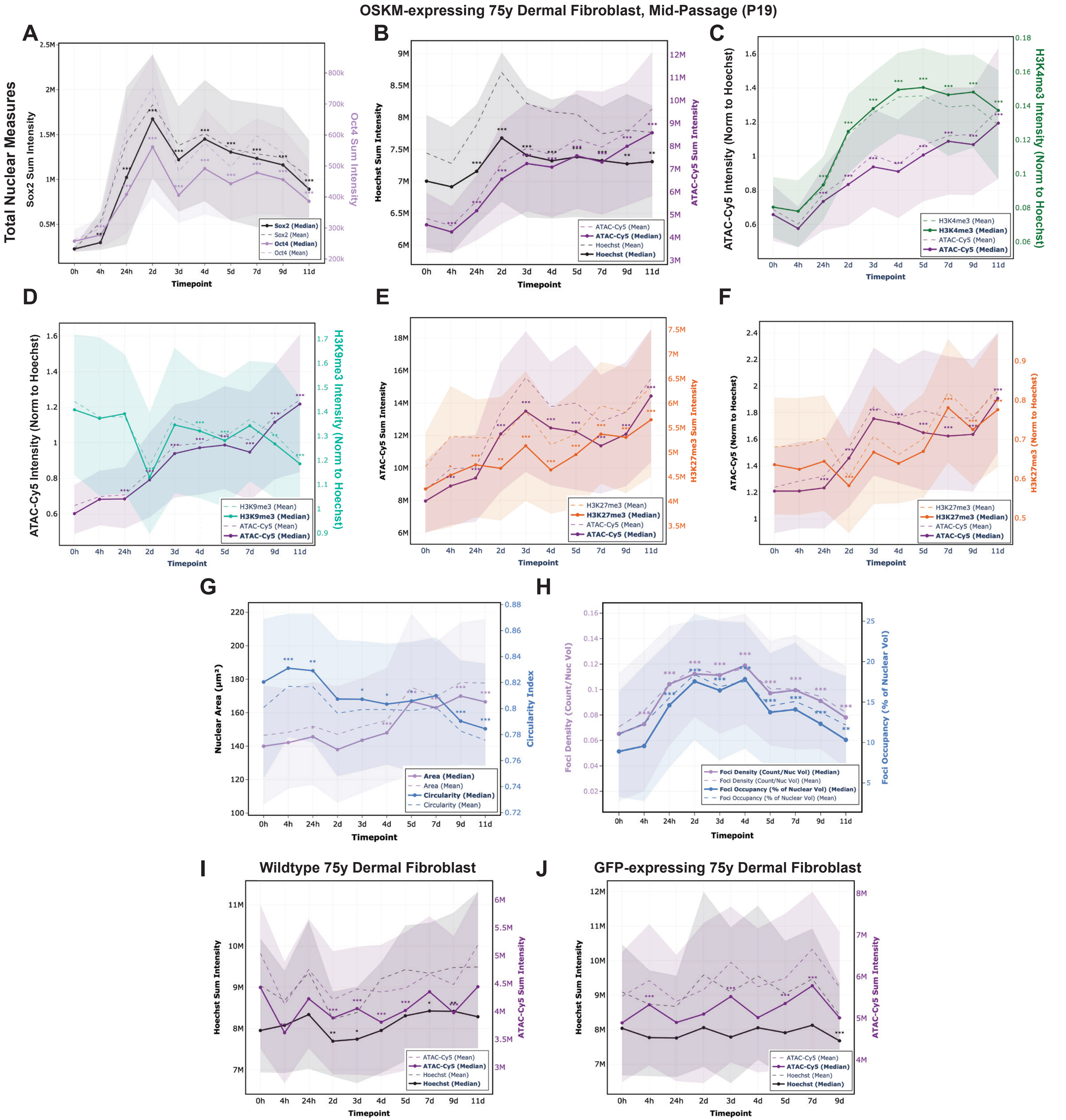
OSKM-expressing 75y Dermal Fibroblast, Mid-Passage (P19)
Total Nuclear Measures
A
Sox2 Sum Intensity
Oct4 Sum Intensity
Sox2 (Median)
Sox2 (Mean)
Oct4 (Median)
Oct4 (Mean)
Timepoint
B
Hoechst Sum Intensity
ATAC-Cy5 Sum Intensity
ATAC-Cy5 (Mean)
ATAC-Cy5 (Median)
Hoechst (Mean)
Hoechst (Median)
C
ATAC-Cy5 Intensity (Norm to Hoechst)
H3K4me3 Intensity (Norm to Hoechst)
H3K4me3 (Mean)
H3K4me3 (Median)
ATAC-Cy5 (Mean)
ATAC-Cy5 (Median)
D
ATAC-Cy5 Intensity (Norm to Hoechst)
H3K9me3 Intensity (Norm to Hoechst)
H3K9me3 (Mean)
H3K9me3 (Median)
ATAC-Cy5 (Mean)
ATAC-Cy5 (Median)
E
ATAC-Cy5 Sum Intensity
H3K27me3 Sum Intensity
H3K27me3 (Mean)
H3K27me3 (Median)
ATAC-Cy5 (Mean)
ATAC-Cy5 (Median)
F
ATAC-Cy5 (Norm to Hoechst)
H3K27me3 (Norm to Hoechst)
H3K27me3 (Mean)
H3K27me3 (Median)
ATAC-Cy5 (Mean)
ATAC-Cy5 (Median)
G
Nuclear Area (µm²)
Circularity Index
Area (Median)
Area (Mean)
Circularity (Median)
Circularity (Mean)
H
Foci Density (Count/Nuc Vol)
Foci Occupancy (% of Nuclear Vol)
Foci Density (Count/Nuc Vol) (Median)
Foci Density (Count/Nuc Vol) (Mean)
Foci Occupancy (% of Nuclear Vol) (Median)
Foci Occupancy (% of Nuclear Vol) (Mean)
I
Wildtype 75y Dermal Fibroblast
Hoechst Sum Intensity
ATAC-Cy5 Sum Intensity
ATAC-Cy5 (Mean)
ATAC-Cy5 (Median)
Hoechst (Mean)
Hoechst (Median)
J
GFP-expressing 75y Dermal Fibroblast
Hoechst Sum Intensity
ATAC-Cy5 Sum Intensity
ATAC-Cy5 (Mean)
ATAC-Cy5 (Median)
Hoechst (Mean)
Hoechst (Median)
0h 4h 24h 2d 3d 4d 5d 7d 9d 11d

**Figure S5 | Transcription factor kinetics, epigenetic trajectories, and control baseline metrics during OSKM reprogramming.** (A) Expression kinetics of integrated pluripotency factors during the reprogramming time-course, quantified via Sox2 and Oct4 Sum Intensities. (B) Reprogramming accessibility dynamics tracked by ATAC-Cy5 Sum and Hoechst Sum Intensities. (C–F) Multiplexed immunofluorescence time-course trajectories plotted alongside ATAC-Cy5 intensities to map histone modifications with global chromatin accessibility. Graphs display H3K4me3 Intensity (C) and H3K9me3 Intensity (D) normalized to Hoechst, as well as raw H3K27me3 Sum Intensity (E) and H3K27me3 normalized to Hoechst (F). (G–H) Extended global morphological and spatial subnuclear metrics, detailing temporal shifts in Nuclear Area and Circularity Index (G), and Foci Density and Foci Occupancy as a percentage of nuclear volume (H). (I–J) Negative control time-courses tracking ATAC-Cy5 and Hoechst Sum Intensities in matched wildtype 75-year-old dermal fibroblasts (I) and GFP-expressing 75-year-old dermal fibroblasts (J). For all line graphs, shaded regions represent the interquartile ranges. Data are representative of single 96-well plate experiments. Each time point represents a pool of 3 replicate wells, with a minimum of $n \geq 499$ single nuclei analyzed per condition for Sox/Oct (A); $n \geq 706$ single nuclei analyzed for Hoechst and H3K4me3 (B, C); $n \geq 727$ single nuclei for H3K9me3 (D); $n \geq 458$ single nuclei for H3K27me3 (E, F); $n \geq 1553$ single nuclei analyzed for foci across 6 well replicates (G, H); $n > 650$ single nuclei analyzed for wildtype and GFP control experiments across 6 well replicates (I, J). $N \geq 3$ biological replicates. Asterisks denote statistical significance of each indicated timepoint relative to the 0h somatic baseline (*** $P < 0.001$, ** $P < 0.01$, * $P < 0.05$; two-sided Mann-Whitney U tests with Bonferroni correction).

### WT, OSK, OSKM - 75y Dermal Fibroblasts, high passage

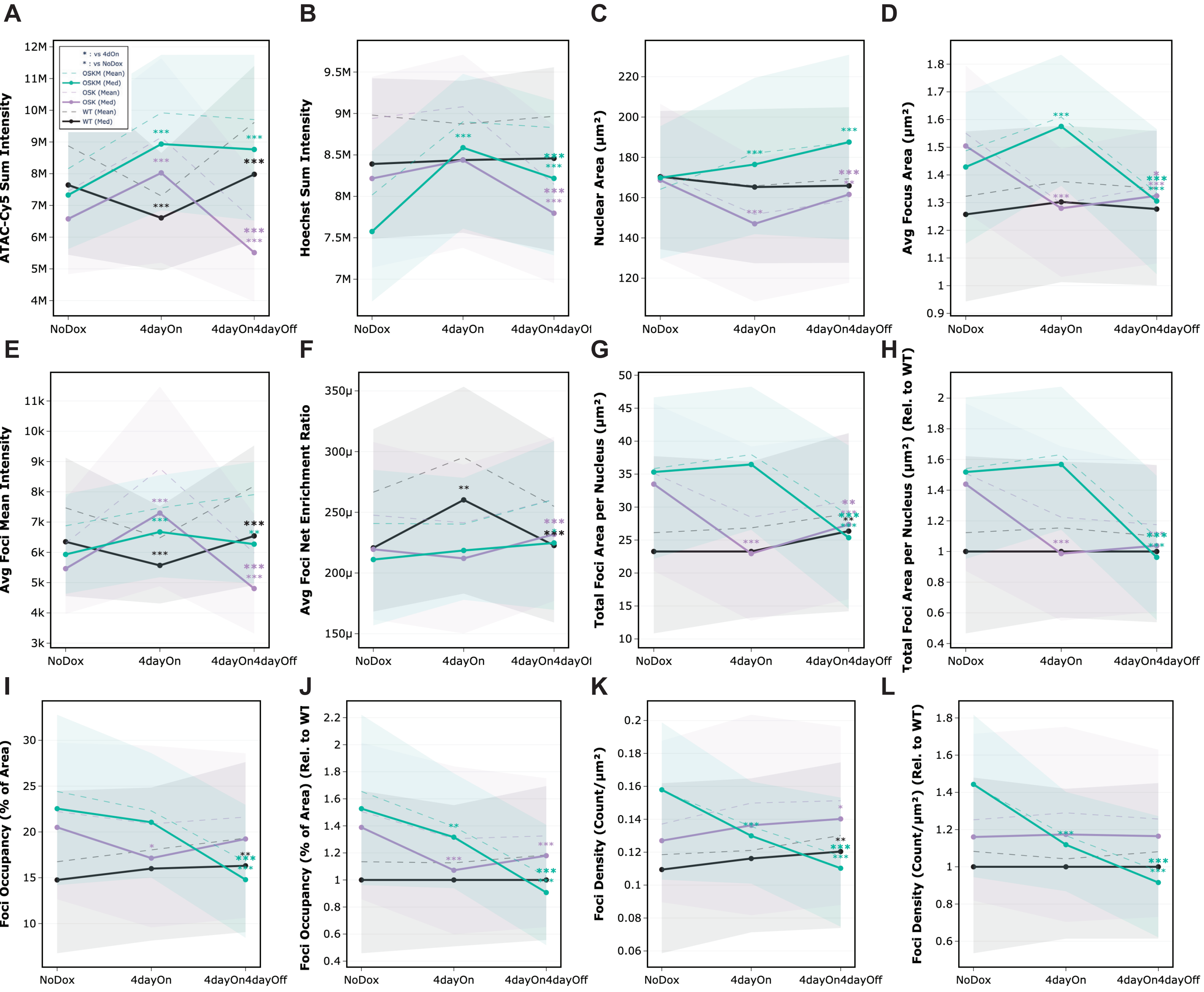


### WT, GFP, OSKM - 75y Dermal Fibroblasts, high passage

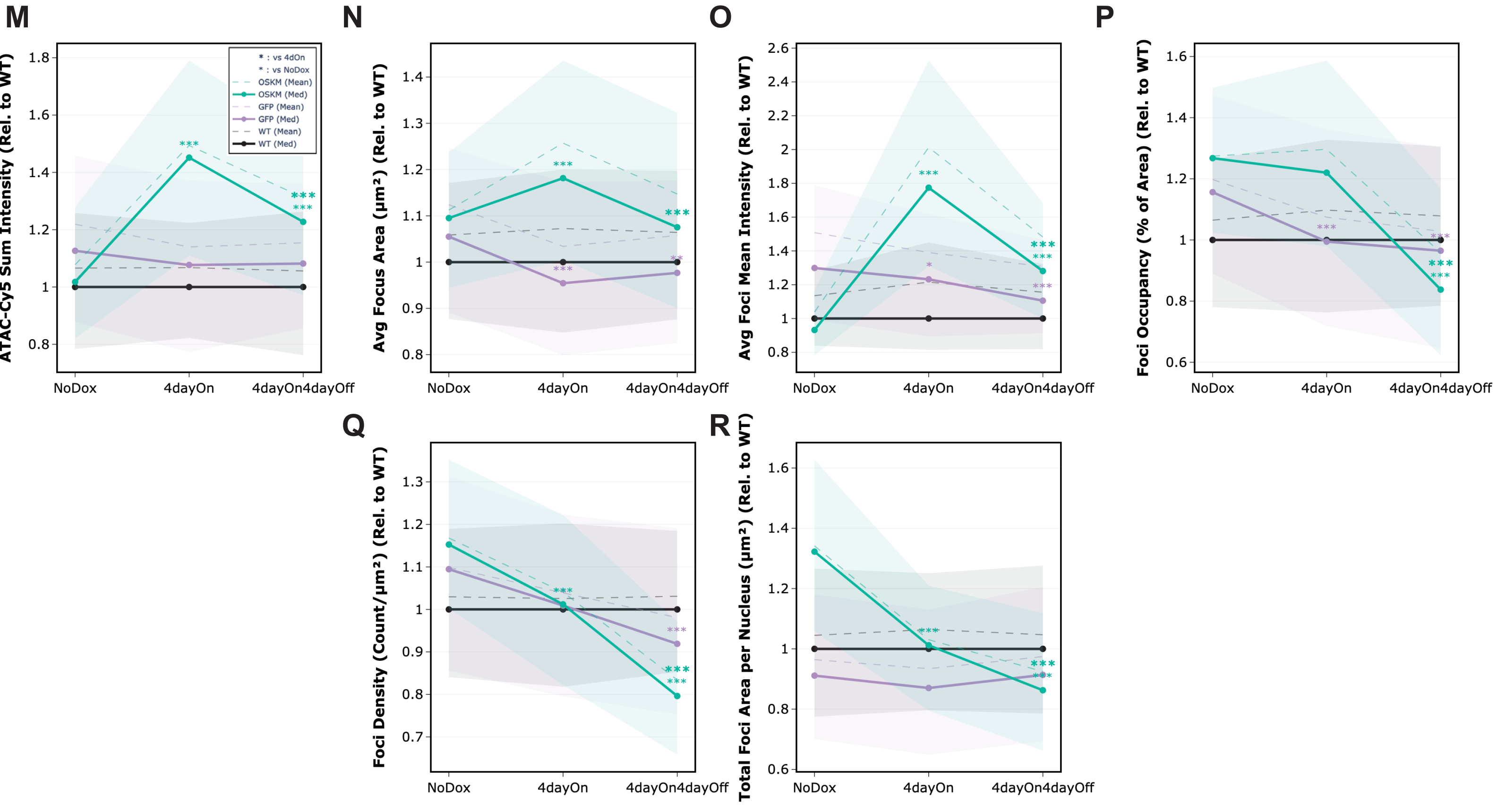

**Figure S6 | Chromatin dynamics for transient OSK and OSKM reprogramming. (A–L)** Extended quantitative metrics for the transient OSK and OSKM induction and recovery experiment in 75-year-old high-passage dermal fibroblasts. Panels include unnormalized ATAC Sum Intensity (A), Hoechst Sum Intensity (B), Nuclear Area (C), Average Focus Area (D), Average Focus Net Enrichment Ratio (E), Total Foci Area per Nucleus (F), and Foci Density (K). Corresponding metrics normalized relative to the matched wildtype (WT) baseline are displayed for Total Foci Area per Nucleus (H), Foci Occupancy as a percentage of nuclear area (J), and Foci Density (L). (M–R) Quantitative metrics from an independent transient reprogramming cohort comparing OSKM induction to a matched GFP-expressing control line. Line graphs display metrics normalized relative to the WT baseline, including ATAC Sum Intensity (M), Average Focus Area (N), Average Focus Raw ATAC Mean Intensity (O), Foci Occupancy as a percentage of nuclear area (P), Foci Density (Q), and Total Foci Area per Nucleus (R). For all line graphs, solid and dashed lines represent the median and mean values, respectively, with shaded regions denoting the interquartile ranges. Bold asterisks indicate statistical significance of the treatment condition at a given timepoint relative to the control (No Dox), non-bold asterisks represent significance between 4 day On and Recovery (*** $P < 0.001$, ** $P < 0.01$, * $P < 0.05$; two-sided Mann-Whitney U tests with Bonferroni correction).

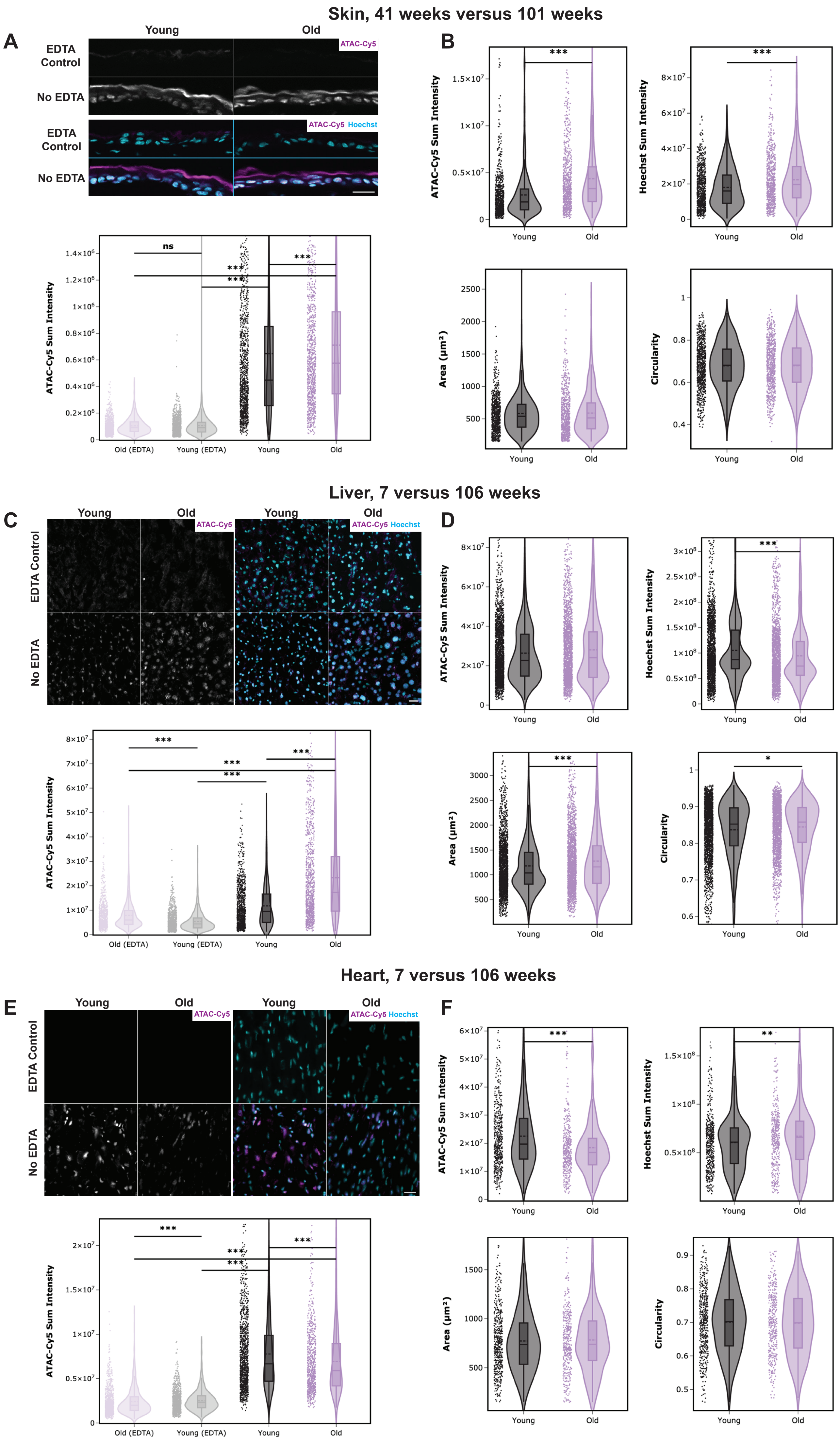
Skin, 41 weeks versus 101 weeks
A
Young
Old
EDTA Control
No EDTA
ATAC-Cy5
ATAC-Cy5 Hoechst
ATAC-Cy5 Sum Intensity
ns
***
Old (EDTA)
Young (EDTA)
Young
Old
B
ATAC-Cy5 Sum Intensity
Hoechst Sum Intensity
Area (µm²)
Circularity
Liver, 7 versus 106 weeks
C
D
*
Heart, 7 versus 106 weeks
E
F
**

**Figure S7 | *In situ* ATAC-see specificity validation, unnormalized intensity distributions and morphological metrics.** (A,C,E) Representative images of ATAC-Cy5 (magenta) and merged channels (ATAC-Cy5, magenta and Hoechst, cyan) in the presence and absence of EDTA, for young and old samples, displayed alongside corresponding violin plots of ATAC-Cy5 Sum Intensity. The baseline autofluorescence and non-specific signal in EDTA controls is significantly lower than true samples, demonstrating the assay specificity. Data presented represents a single biological replicate, and was confirmed across more than two biological replicates to ensure reproducibility. Scale bars, 20μm. Skin (A, B; 41 weeks vs. 106 weeks), Liver (C, D; 7 weeks vs. 103 weeks) and Heart (E, F; 7 weeks vs. 103 weeks). Asterisks denote statistical significance across the single-nuclear populations (*** $P < 0.001$; two-sided Mann-Whitney U tests with Bonferroni correction). (B, D, F) Comprehensive single nuclear distributions of raw, unnormalized metrics across an additional, independent *in vivo* aging cohort. Panels detail ATAC-Cy5 Sum Intensity, Hoechst Sum Intensity, Nuclear area ($\mu m^2$) and Nuclear Circularity for Skin (B), Liver (D) and Heart (F). Data represents a single biological replicate for $N$ = 6 (Skin), 4 (Liver) and 7 (Heart). Asterisks denote statistical significance across the individual nuclear populations (*** $P < 0.001$, ** $P < 0.01$, * $P < 0.05$; two-sided Mann-Whitney U tests).
